# The multimerization pathway of the glucocorticoid receptor

**DOI:** 10.1101/2024.12.12.628195

**Authors:** Andrea Alegre-Martí, Alba Jiménez-Panizo, Agustina L. Lafuente, Thomas A. Johnson, Inés Montoya-Novoa, Montserrat Abella, Paloma Pérez, Juan Fernández-Recio, Diego M. Presman, Gordon L. Hager, Pablo Fuentes-Prior, Eva Estébanez-Perpiñá

## Abstract

The glucocorticoid receptor (GR) is a leading drug target due to its anti-inflammatory and immunosuppressive roles. The functional oligomeric conformation of full-length GR (FL-GR), which is key for its biological activity, remains disputed. Here we present a new crystal structure of agonist-bound GR ligand-binding domain (GR-LBD) comprising eight copies of a non-canonical dimer. The biological relevance of this dimer for receptor multimerization in living cells has been verified by studying single-and double-point mutants of FL-GR in fluorescence microscopy (Number & Brightness) and transcriptomic analysis. Self-association of this GR-LBD basic dimer in two mutually exclusive assemblies reveals clues for FL-GR multimerization and activity in cells. We propose a model for the structure of multidomain GR based on our new data and suggest a detailed oligomerization pathway. This model reconciles all currently available structural and functional information and provides a more comprehensive understanding of the rare glucocorticoid resistance disorder (Chrousos syndrome).

## Introduction

The glucocorticoid receptor (GR/NR3C1) is a ligand-activated transcription factor essential for life (1–3). GR belongs to the steroid receptor subfamily of nuclear receptors (NRs) that also includes the mineralocorticoid receptor (MR/NR3C2), the progesterone receptor (PR/NR3C3), the androgen receptor (AR/NR3C4), as well as estrogen receptors α and β (ERα/NR3A1 and ERβ/NR3A2) (4–6). Full-length GR (FL-GR) shares a modular architecture with the other steroid receptors that is accountable for their (patho-)physiological actions: an intrinsically-disordered N-terminal domain (NTD), a highly conserved DNA-binding domain (DBD), a flexible hinge, and a mostly helical ligand-binding domain (LBD) (Figure 1A) (4, 6). GR and related oxosteroid receptors (AR, PR, and MR) display a C-terminal extension named F-domain (Figure 1A) (6). GR binds glucocorticoids (GCs; either endogenous stress hormones such as cortisol, or synthetic, e.g., dexamethasone (DEX)) in an internal ligand-binding pocket (LBP) (4, 7–9). In addition, the LBD displays two solvent-exposed protein-protein binding platforms: activation function-2 (AF-2) and binding function-3 (BF-3), which are responsible for the recruitment of coregulators and the chaperone/cochaperone complex (Hsp90-p23) (10–16).

**Figure 1 Legend.**
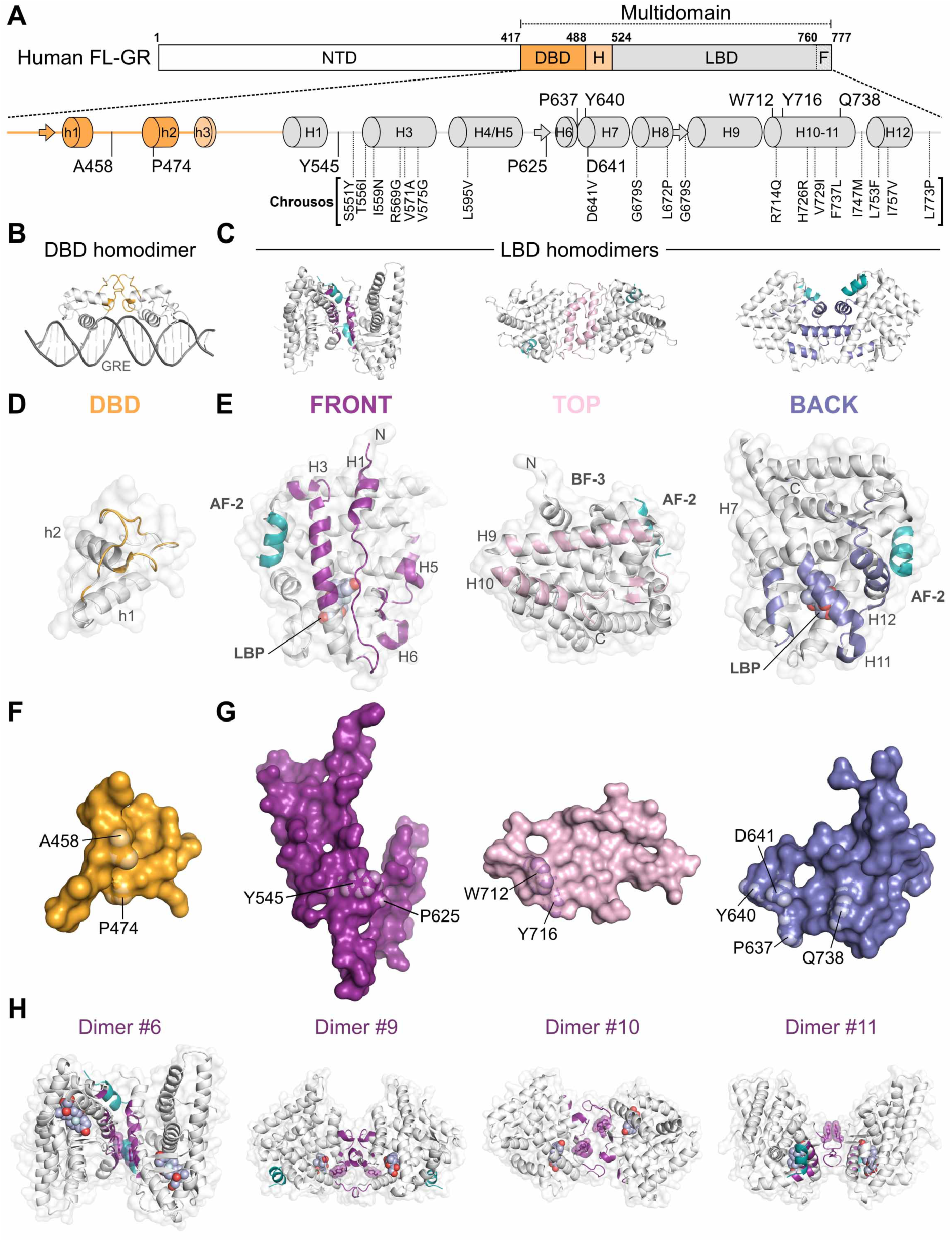
Both DBD and LBD modules of the GR are important for homodimerization. (A) Top panel: Schematic representation of FL-GR domain organization. The NTD is followed by the ‘core’ or multidomain GR, featuring the DBD, the hinge, and the LBD. Lower panel: secondary structure of multidomain GR, highlighting residues important for receptor multimerization. Missense mutations or SNPs linked to Chrousos syndrome are also indicated; note that they cluster in the L1-3/H3 stretch and the most C-terminal helices, H10/H11 and H12. (B-G) Summary of current structural information on the dimerization of individual DBD and LBD modules. (B) Structure of the canonical GR-DBD homodimer bound to the GRE from the hypersensitive gene, *SGK1* (PDB entry 7PRW). Monomers are shown as gray cartoons with secondary structure elements involved in interdomain interactions colored yellow. (C) Representative structures of symmetric GR-LBD homodimers corresponding to the previously described front-to-front, top-to-top, and back-to-back arrangements (PDB 7YXP, 4P6W, and 4LSJ, respectively). For simplicity, no example of base-to-base dimers is included here. Monomers are shown as gray cartoons and their interacting secondary structure elements are colored violet, pale pink or blue, respectively. Coregulator peptides bound to the AF-2 cleft are shown as teal cartoon. (D-E) One monomer from each of the dimers shown in panels B-C is rotated 90° to present the dimerization interface to the viewer. Monomers are depicted as cartoons with semi-transparent surfaces superimposed. Major secondary structure elements are colored as above and labeled. The DEX molecule bound in the LBP is represented as light-blue spheres with red-colored oxygen atoms. (F, G) Close-ups of the dimerization interfaces, shown as solid surfaces color-coded as in panels B-E. The location of selected residues important for multimerization is indicated. (H) Out of 20 distinct homodimeric arrangements of GR-LBD included in our recently presented catalog, four non-canonical GR-LBD homodimers appear to correspond to transcriptionally active states of the receptor in cells. These GR-LBD dimers -arbitrarily numbered #6, #9, #10 and #11-are depicted as cartoons with overlaid semi-transparent surfaces. The interacting areas are colored as above. Residues Tyr545/Tyr545’ are highlighted.

*NR3C1* is constitutively expressed in almost all human cells, where it mediates GCs actions (1, 17, 18). GR transcriptional activity requires binding to GC response elements (GRE) in chromatin (Figure 1B) (2, 3, 19–22). Both DNA and hormone act as allosteric GR modulators that regulate receptor oligomerization (7, 23). Ligand binding induces dimerization and docking to the DNA triggers further structural rearrangements that promote the formation of higher-order oligomers, predominantly tetramers (24), through surfaces that are yet to be identified. Other oxosteroid receptors have been shown to oligomerize before DNA binding (24–26). These and other observations suggest a multi-step, highly cooperative oligomerization mechanism. Both the oligomerization state of steroidal receptors, the dynamics of their interaction with chromatin and the active oligomeric conformation of the DNA-bound receptors remain areas of intense research (27–35). Mutations in *NR3C1* are linked to human pathology, leading to either primary generalized GC resistance (PGGR, also known as Chrousos syndrome(36–39)) (Figure 1A) or hypersensitivity (40, 41). GR is one of the major pharmacological targets because of its potent anti-inflammatory and immunosuppressive actions, but patients on chronic GCs develop resistance and serious side effects (42).

Structural biology has been essential to develop therapeutic GR modulators for several pathologies (e.g., asthma, psoriasis, rheumatoid arthritis, cancer, diabetes) or to prevent organ rejection (1, 42–48). The GR-DBD homodimer has been characterized both in the absence and presence of DNA (Figures 1B, D, and F) (49–51). At the same time, numerous crystal structures of isolated GR-LBD bound to agonists or antagonists have been reported (Figures 1C, E, and G) (4, 52, 53). Recently, the structure of multidomain GR (DBD-hinge-LBD) in complex with ligand, DNA, and co-regulator peptide has been solved to medium resolution (54). However, the hinge regions in this structure were not defined by electron density complicating the interpretation of intra-and inter-monomer contacts. Furthermore, no structure of FL-GR has been solved to date and technical limitations to assess the state of the receptor *in vivo* have stimulated intense controversy over the years. Thus, several key questions about GR tertiary and quaternary structure remain open including how disease-linked mutations or post-translational modifications (PTMs) affect receptor architecture (1, 6, 55, 56). We have recently shown that GR-LBD modules can form various quaternary arrangements depending on bound agonists/antagonists and other biochemical parameters. We identified four GR-LBD oligomerization interfaces (termed front, top, back, and base) (Figures 1C, E, and G), the combination of which yielded a catalog of 20 distinct homodimers (52). Furthermore, our results suggested that four topologically distinct, non-canonical GR homodimers could correspond to the transcriptionally active receptor (Figure 1H). On the other hand, our analysis of the Chrousos syndrome-linked mutation, p.Asp641Val, revealed transcriptionally compromised higher-order GR multimers, demonstrating the complexity of GR multimerization (52). Despite these new insights, the precise pathways by which GR monomers associate to form dimers and then tetramers and/or higher-order oligomers remain unclear (24, 26).

Here we present a new crystal structure of agonist-bound GR-LBD featuring an unusually large unit cell and a very high solvent content. The building block of the current structure is a homodimer whose biological relevance has been verified using a battery of relevant FL-GR mutants in fluorescence microscopy (Number and Brightness, N&B) experiments in living cells and RNA sequencing (RNA-seq) analysis. Several mutually exclusive assemblies of this basic dimer in the crystal highlight the versatility of GR-LBD for self-association with implications for FL-GR function in cells. We propose an alternative model of multidomain GR based on this data and suggest pathways for receptor multimerization *in vivo*.

## Materials and methods

### Crystallization and structure determination

Recombinant ancGR2-LBD (corresponding to residues 529–777 of the human receptor) was expressed in *E. coli* in the presence of 50 mM DEX and purified to homogeneity as previously described (7, 52). Concentrated DEX-bound ancGR2-LBD was combined with a 3-fold molar excess of a peptide corresponding to residues Gln18-Lys27 of the small heterodimer partner (SHP/NR0B2, box 1 motif; NH_2_-QGAASRPAILYALLSSSLK-OH; Pepmic) and incubated for one hour at RT.

Drops of the ancGR2-LBD-SHP mixture were equilibrated at 20 °C against 0.085 M sodium cacodylate trihydrate, pH 6.5, 0.17 M ammonium sulfate, 25.5% (w/v) PEG 8000 and 15% (v/v) glycerol, using the sitting drop vapor-diffusion method.

Diffraction data were collected at 100 K at the ALBA-CELLS synchrotron and processed using programs of the CCP4 suite (http://www.ccp4.ac.uk/). The crystal structure was solved and refined using MOLREP, REFMAC5 and COOT from the CCP4 package. Crystal packing was analyzed using PISA (http://www.ebi.ac.uk/), model quality was assessed with MolProbity (http://molprobity.biochem.duke.edu/) and ModFOLD (https://www.reading.ac.uk/bioinf/ModFOLD/). Figures and videos were prepared with PyMOL (http://www.pymol.org).

### Cell line generation and culture

Mammary adenocarcinoma 3617-derived GR knock-out (GR-KO) cells has been previously described (97, 100). Cells were grown in Dulbecco’s modified Eagle’s medium (Invitrogen) supplemented with 5 µg/ml tetracycline (Sigma), 10% fetal bovine serum (FBS, Sigma), sodium pyruvate, non-essential amino acids, and 2 mM L-glutamine maintained in a humidifier at 37 °C. The FBS-supplemented medium was replaced with medium supplemented with charcoal/dextran-treated FBS to remove GCs from cells for 24 h prior to hormone treatment (100 nM DEX for 1h). For N&B assays, cells were transiently transfected with the following variants of GFP-tagged mouse GR: WT, Y551E, P631A, P481/P631A, P643C, Y646A, Y646A/Y722S, W718E, W718S/Y722S, and Q744A, which correspond to human mutations Y545E, P625A, P474R/P625A, P637C, Y640A, Y640A/Y716S, W712E, W712S/Y716S, and Q738A, respectively, using jetOPTIMUS™ or JetPrime reagent (PolyPlus) according to the manufacturer’s instructions. The cell lines used for total RNA-seq experiments were developed in the same GR-KO cell line as previously described (97, 100). Briefly, GFP-tagged variants of mouse GR (WT, P631A, P643A, W718E and W718S/Y722S) were stably integrated into the GT-Rosa26 locus via CRISPR/Cas9 homology-directed repair with puromycin selection and fluorescent-activated cell sorting. GFP-GR variants for RNA-seq and N&B were generated with the QuikChange II XL Site-Directed Mutagenesis Kit (Stratagene) or by Gene Universal.

### N&B experiments and analysis

Images were taken either at the CCR, LRBGE Optical Microscopy Core facility (Bethesda, MD, USA) or at the Weber’s Advance Microscopy Core (Buenos Aires, Argentina) using LSM 780 or LSM 980 laser scanning microscopes (Carl Zeiss, Inc.), respectively, both equipped with an environmental chamber. Cells were excited with a multi-line Argon laser tuned at 488 nm and imaged for 20-120 min after DEX addition using a 63× oil immersion objective (NA = 1.4). Fluorescence was detected with a GaAsP detector in photon-counting mode.

N&B measurements were done as previously described (24, 25). For each studied cell a single-plane stack of 120 images (256 × 256 pixels) were taken with a pixel size of 80 nm and pixel dwell times of 6.27 and 8.19 µs in the LSM 780 and LSM 980 scopes, respectively. In all stacks, we discarded the first 5 images to reduce the effect of photobleaching. The frame time under these conditions is 0.97 s (LSM 780) or 1.26 s (LSM 980), which guarantees independent sampling of molecules according to previously reported fluorescence correlation spectroscopy (FCS) measurements (101). Each stack was further analyzed using the N&B routine of the SimFCS 2.0 software (Global Dynamics), in which the average fluorescence intensity () and its variance (σ2) at each pixel of an image are determined from the intensity values obtained at the given pixel along the image stack. The apparent brightness (B) is calculated as the ratio of σ2 to while the apparent number of moving particles (N) corresponds to the ratio of to B (102). We used the SimFCS 2.0 software to classify pixels as occurring in the nucleus or the MMTV array (103) according to their intensity values.

Cells were selected for analysis following these criteria: (i) in the case of stimulated cells, an accumulation of signal at the array must be visible, (ii) the average apparent number of molecules (N) in the nuclear compartment must have a range of 3–30 units in all cases, (iii) no saturation of the detector at any pixel (N < 60), and (iv) bleaching cannot exceed 5–10%. In previous work it was demonstrated that B is equal to the real brightness ε of the particles plus one.(102) Therefore, ε at every pixel of images can be easily extracted from B measurements. Importantly, this analysis only provides information regarding the moving or fluctuating fluorescent molecules since fixed molecules (relative to our frame time) will give B values equal to 1. The experiments were independently repeated at least two times for each treatment/condition.

### Total RNA collection and sequencing

RNA was isolated using the PureLink RNA kit (Thermo Fisher Scientific) following the manufacturer’s instructions. We collected three biological replicates of each condition and assessed sample quality using the Agilent Bioanalyzer. Strand-specific sequencing libraries were generated from rRNA-depleted (Illumina RS-122-2301) total-RNA samples, using Illumina Stranded Total RNA (Illumina20020596) according to the manufacturer’s instructions. Raw reads were demultiplexed into Fastq format allowing up to one mismatch using Illumina Bcl2fastq v2.17. Reads of the samples were trimmed for adapters and low-quality bases using Cutadapt1.18. RNA-seq alignment to the mouse mm10 genome was performed with STAR (104). DESEQ2 (105) was used to normalize the data by read depth, identify differentially expressed genes for each form of GR (DEX/vehicle), and calculate log2 fold changes (FC) and false discovery rates (FDR) for each gene.

### Docking and other bioinformatics analyses

Browser was used for superimposing structures and RMSD calculation (Molsoft LLC). Solvent-protected areas were calculated with PISA (http://www.ebi.ac.uk/). Energy terms were calculated with the bindEy module of pyDock (70).

We explored the multimerization potential of multidomain GR using pyDock docking and scoring method (70). First, protein molecules were prepared by removing all cofactors and heteroatoms, and missing side chains were modeled with SCWRL 3.0 (106). Then, the Fast Fourier Transform (FFT)-based docking programs FTDock(107) (with electrostatics and 0.7-Å grid resolution) and ZDOCK 2.1 (108) were used to generate 10,000 and 2,000 rigid-body docking poses, respectively. These were merged in a single pool for subsequent pyDock scoring, based on energy terms previously optimized for rigid-body docking. The pyDock binding energy is basically composed of accessible surface area-based desolvation, Coulombic electrostatics, and VdW energy terms. Electrostatics and VdW contributions were limited to –1.0/+1.0 and 1.0 kcal/mol for each inter-atomic energy value, respectively, to avoid excessive penalization from possible clashes derived from the rigid-body approach.

From the resulting docking poses, NIP values were obtained for each residue with the built-in patch module of pyDock, implementing the pyDockNIP algorithm (109). A NIP value of 1 indicates that the corresponding residue is involved in all predicted interfaces of the 100 lowest energy docking solutions, while a value of 0 means that it appears as expected by random chance. Finally, a negative NIP value implies that the residue appears at the low-energy docking interfaces less often than expected by random chance. Usually, residues with NIP ≥ 0.2 are considered as part of a hot-spot when using FTDock.

## Results

### Tyr545-centered, parallel homodimers form the basic repeat unit of the new GR-LBD crystals

Using the resurrected variant of human GR, ancGR2,(7) which shows higher solubility and stability than ‘modern’ GR, we obtained a new crystal form of GR-LBD characterized by a large unit cell (orthorhombic space group P2_1_2_1_2 with a = 264.4, b = 265.5, c = 109.7 Å, α = β = γ = 90°), together with an unusually high solvent content (≈70%), with channels that are up to 40 Å wide (Figures 2A-B). As a result, the asymmetric unit (ASU) of the crystals contains 16 GR-LBD monomers, which have been alphabetically labeled from A to P (Figures 2C and S1). This is in contrast with the 22-26 copies that would have been expected in protein crystals with the typical solvent content of ≈50% (see Supplementary Table S1 for a summary of data collection and refinement statistics). The high solvent content of the crystals and the sparse number of GR-LBD molecules present in the crystal lattice result in a more physiological setting than previously reported GR-LBD crystals, as discussed below. The 16 crystallographically independent monomers can be organized into two different settings of the ASU (Figures 2A-C). These two settings will be examined in detail below in connection with the identification of biologically relevant interfaces.

**Figure 2 Legend.**
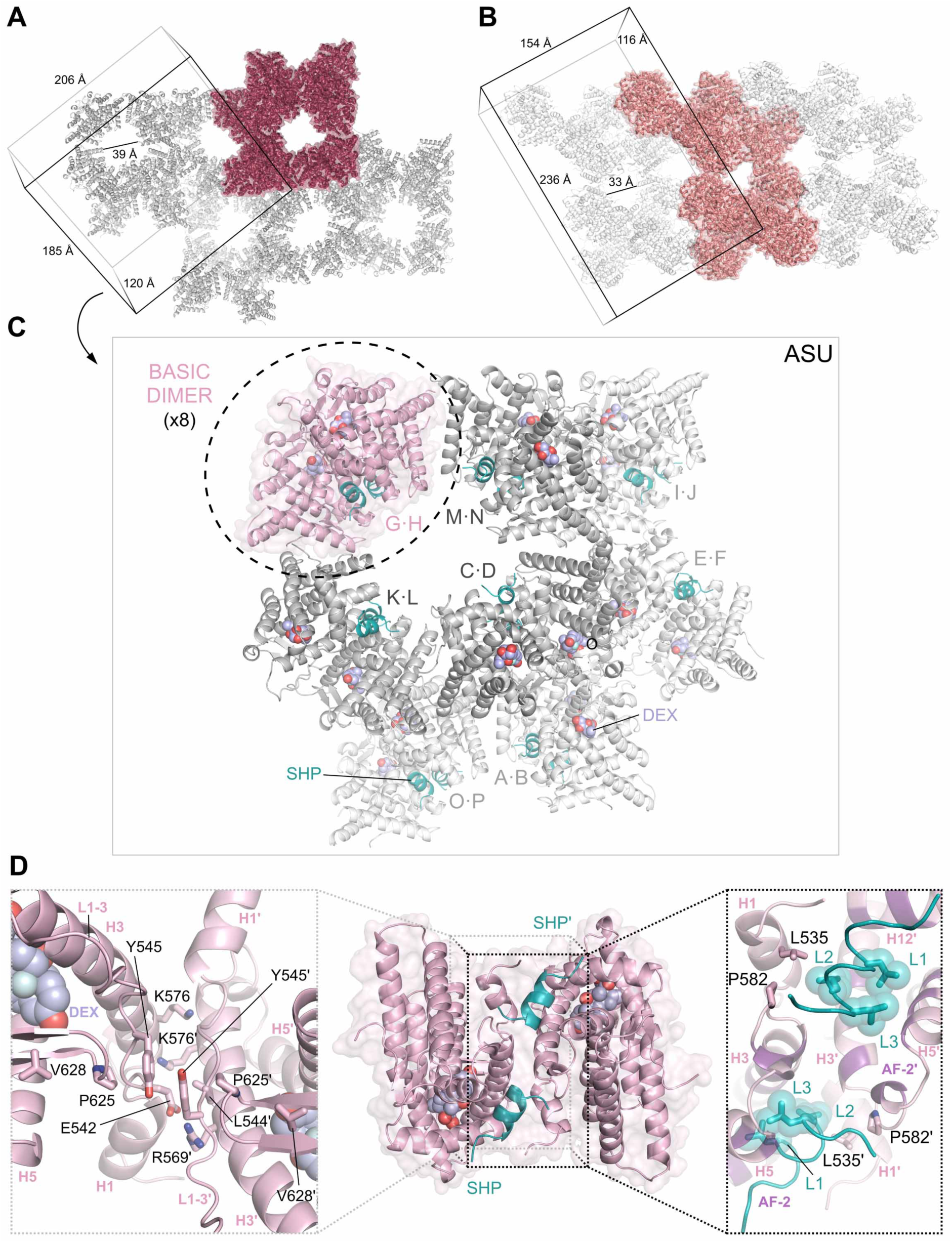
A novel crystal structure of GR-LBD reveals the structural determinants of GR multimerization. Crystals feature an unusually high solvent content in addition to a large unit cell of about 7.7 × 10^6^ Å^3^, with large and wide channels filled with solvent molecules. Panels A and B depict the two different ways of choosing the ASU of the current crystals that maximize the number and quality of inter-monomer contacts. These two arrangements of the 16 independent GR-LBD monomers likely capture (patho)physiologically relevant protein-protein contacts *in vivo* (see main text for details). (A) Choice of the ASU centered on residues of helices H9/H10 and the connecting loop (L9-10). Because of the prominent role of the Trp712 indole rings in this arrangement, it will be referred to as the ‘Trp712 setting’. The 16 crystallographically independent GR-LBD monomers are colored in burgundy. (See Figure 5 for details of inter-monomer contacts in this setting). (B) An alternative choice of the ASU will be referred to as the ‘Pro637 setting’ as the major organizing elements are residues of the L6-7 loop, most notably Pro637. The 16 independent GR-LBD monomers are colored in coral. (See Supplementary Figure 4 for details of inter-monomer contacts in this setting). (C) A view of the ASU of the crystals in the Trp712 setting, with all 16 independent GR-LBD molecules represented as cartons. Note that these 16 monomers associate to form eight similar parallel homodimers, which form the basic repeating unit of the ASU. One of these basic dimers is colored pale pink and the other seven dimers are shown in different shades of gray depending on their position along the axis perpendicular to the plane of the page. The SHP coregulator peptides and DEX molecules are color-coded as in Figure 1. (D) The basic GR-LBD homodimer belongs to class #6 in our catalog of possible homodimeric arrangements (Figure 1H and ref. (52)). The dimer is shown as a pale pink cartoon at the center of the panel, with coregulator peptides and DEX molecules colored as in C. Close-ups of the protein-protein dimer interface are shown to the right and left. Note the central position occupied by the stacked phenol rings of residues Tyr545/Tyr545’, as well as the stabilizing role of Pro625 and Val628 (left panel). (Residues from the second monomer in the basic dimers are primed). Interactions of SHP residues Leu21, Leu25 and Leu26 with the AF-2 cleft are highlighted in the right panel. (Labeled L1, L2 and L3 in the figure for simplicity). Note that the bound peptides cross-connect the two monomers: in addition to occupying the AF-2 cleft of its ‘own’ LBD in a canonical manner, the more C-terminal SHP residues contact the neighboring GR module.

Important differences in the main chain path of the 16 independent monomers are limited to two zones located on opposite poles of the domain: the loop connecting helices H9 and H10 (L9-10) together with the N-terminal end of H10, and a surface formed by the neighboring L1-3, L6-7 and L11-12 loops (Figure S1A). Both main and side chains of all monomers are well defined by electron density, with the only exception of monomer H in which side chain atoms are much more poorly defined due to enhanced thermal motion (Figures S1B-D and S2A-B).

Most notably, the L9-10 loop adopts in several monomers a ‘closed’ conformation characterized by a Glu705-Gly706 tight turn with a solvent-pointing Glu705 side chain and H10 N-terminally extended to residue Asn707 (Figure S1E). We use throughout the manuscript a numbering system based on the canonical human sequence, UniProt entry P04150-1, corresponding to variant GRα-A. This doubly capped structure of H10 is similar to the path followed in the high-resolution structure of mouse GR-LBD featuring mutations Val702Ala and Glu705Gly, which lead to enhanced stability of the LBD module (Protein Data Bank (PDB) 3MNP (57)). By contrast, residue Glu705 points towards the interhelical space in other monomers and the L9-10 loop adopts an ‘open’ conformation that is not well defined by electron density in most monomers, indicating enhanced mobility (Figures S1D and S2A-C). Interestingly, all but one dimer (O·P) feature GR-LBD modules in which one loop L9-10 adopts the closed, while the other is found in the open conformation. This observation suggests that the flip-flop movement of residue Glu705 and the accompanying rearrangements in this region are linked to L1-3 and thus to homodimer formation, in line with the results of Seitz and coworkers (57) and previous molecular dynamics simulations (58).

The 16 copies of the GR-LBD monomers are organized into eight similar, parallel homodimers that belong to class #6 as listed in our recently presented catalog of possible homodimeric arrangements (Figures 1H and 2C-D) (52). These ‘basic’ homodimers are generated by contacts between solvent-exposed residues in loops L1-3 and helices H3 from the two monomers (Figure 2D). These residues are strictly conserved or conservatively replaced from fishes to humans, particularly in human GR, highlighting the biological relevance of the arrangement. At the center of this interface, the phenolic side chains of residues Tyr545/Tyr545’ stack on each other and are perfectly defined by electron density in all pairs (Figures 2D and S1F). This is in addition to strong salt bridges between the carboxylate of Glu542 and the guanidinium group of Arg569’ and *vice versa*, which are well protected from bulk solvent by the aliphatic side chains of Leu544’/544. Disruption of these salt bridges might explain the functional impairment of the recently reported mutation linked to Chrousos syndrome, p.Arg569Gln (Supplementary Table S2) (59). Further, the aliphatic parts of residues Lys576/576’ make Van-der-Waals (VdW) contacts with each other and in addition engage in both VdW and electrostatic interactions with residues Ile539’/Ile539 from the neighboring monomer. Altogether, 565 ± 15 Å^2^ are protected from bulk solvent in these parallel, Tyr545-centered homodimers (see Supplementary Table S3 for the contribution of different energy terms to dimer stability). Of note, the conformation of the Tyr545 side chain is stabilized by strong Van-der-Waals interactions with a residue from the underlying loop connecting H5 and H6, the strictly conserved Pro625 (Figure 2D). Several previous investigations have assessed the relevance of the L5-6 loop, especially of residue Ile628, for GR oligomeric state and activity (24, 26). For simplicity, in the following we will refer to this local structure as the ‘Tyr545 interface’, with the understanding that other neighboring residues of the L1-3/H3 area and the underlying Pro625/Ile628 are integral elements of the interface.

### Tyr545-centered, non-canonical homodimers are essential for GR multimerization and transcriptional activity

The presence of eight similar Tyr545-centered GR-LBD dimers in the current crystals suggested an enhanced stability of this homodimeric arrangement in solution. This observation, along with our previous data showing the impact of the Y545A mutation on GR oligomerization and transcriptional activity (52), prompted us to assess in more detail its relevance in living cells. To this end, we generated a new FL-GR mutant of the central interface residue, Tyr545 (variant GR^Y545E^), aimed at exploiting the electrostatic repulsion of approaching Glu545 side chains (Figure 3A). In addition, we mutated a residue that makes important contacts with the phenol ring of Tyr545, Pro625 (variant GR^P625A^) (Figures 2D and 3A). Next, we studied the oligomerization behavior of these mutants by N&B, both in the nucleoplasm and DNA-bound (Figures 3B and Supplementary S3A), essentially as previously reported (24, 52, 60).

**Figure 3 Legend.**
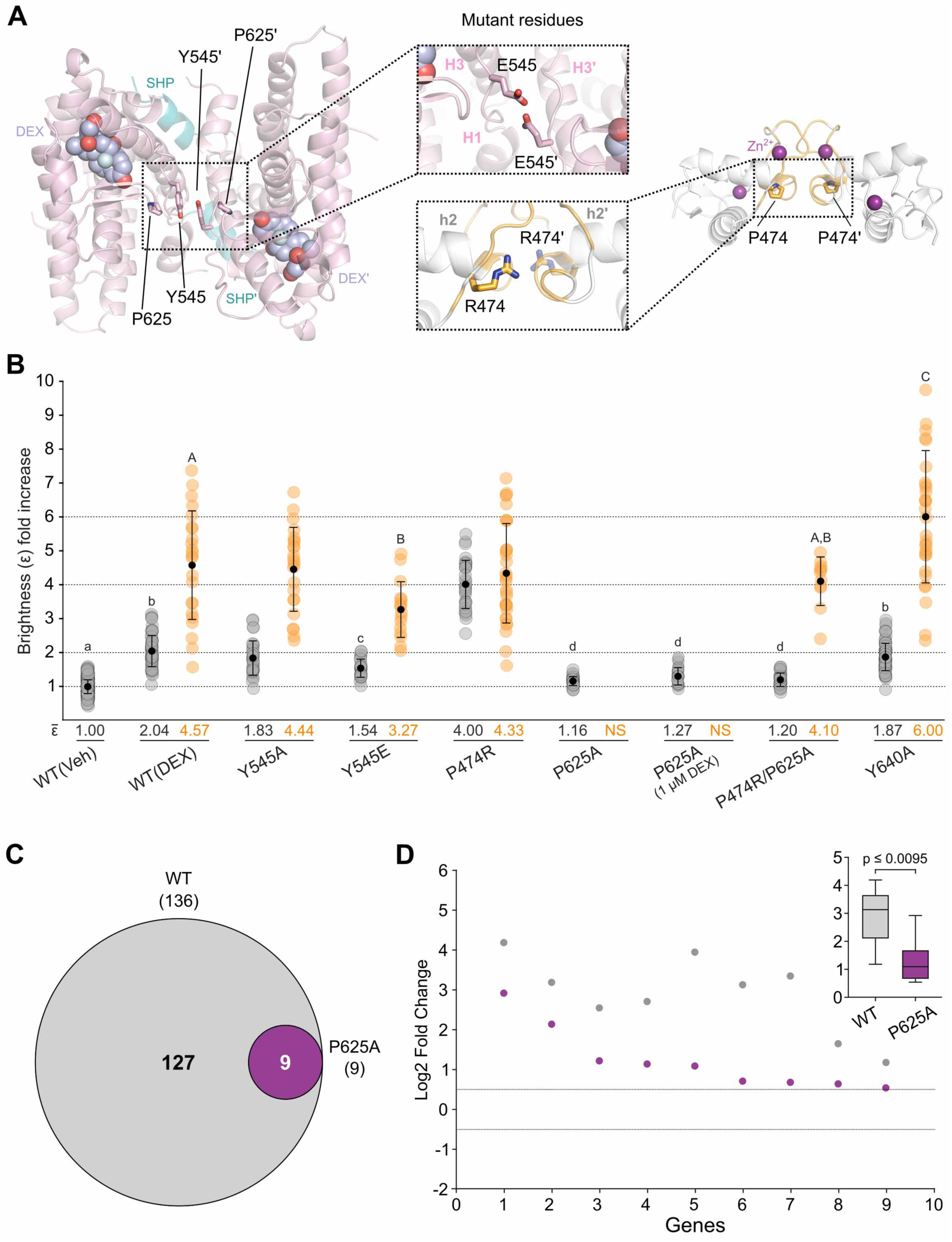
The Tyr545-centered non-canonical homodimer is important for FL-GR dimerization and transcriptional activity. (A) 3D representations of the Tyr545-centered LBD homodimer (to the left) and the canonical DBD homodimer (to the right). The SHP coregulator peptides and DEX moleculas bound the the LBD are color-coded as in Figure 1. Residues mutated in the current investigations are shown as color-coded sticks and labeled. The central panels show close-ups of the models with introduced mutations. Note that the presence of glutamate residues at positions 545/545’ would result in strong electrostatic repulsion at the LBD-LBD’ interface. On the contrary, the replacement of Pro474 in the DBD by a bulkier arginine would allow for important VdW and electrostatic interactions with the neighboring monomer. (B) N&B assay. The fold increase in molecular brightness (ε), is given relative to the value for the non-stimulated WT receptor (vehicle, Veh). The results in the nucleoplasm and at the array are represented as gray-and orange-colored dots, respectively. Results for the GR^Y545A^ and GR^P747R^ variant were taken from our previous publication(52) and are shown here for comparison purposes only. Note that no signal was observed for the GR^P625A^ mutant at the array (NS). The sample mean is shown as a black point in the middle of each dot plot, together with the standard deviation bars. Dot plots with different superscript letters are significantly different from each other (p < 0.05, one-way ANOVA followed by Tukey multiple comparisons test). Statistical analysis of the nucleoplasm and array data were done separately (small and capital letters, respectively). (C) Venn diagram comparing hormone-regulated protein-coding genes (2 h DEX treatment/vehicle) in variant GR^P625A^ and the WT receptor. Three replicates were used per condition. The total number of hormone responsive genes (FDR < 0.01, |Log2 FC| > 0.5) is given in parentheses. (D) Scatter plot of Log2 FC for the nine responsive genes shared between WT GR and GR^P625A^. Dashed horizontal lines denote Log2 FC +/-0.5. Box and whiskers plot of the same data displays interquartile range depicting the 25^th^, 50^th^ and 75^th^ percentile as box with the median as a black bar. Statistical analysis was performed using a two-tailed non-parametric t-test. Note that even the nine genes that were up-regulated by GR^P625A^ were transcribed at significantly lower levels.

Mutants translocated into the nucleus upon cell stimulation with 100 nM DEX (Figure S3A), although GR^P625A^ remained partly cytoplasmic after 40 minutes. Successful nuclear translocation suggests that neither the folding of the receptor nor its interactions with the import machinery were significantly affected. GR^P625A^ was mostly monomeric in the nucleoplasm and could not be detected at the mouse mammary tumor virus (MMTV) array (Figure 3B), consistent with a recent report (61). A 10-fold higher DEX concentration was barely able to increase the dimerization of this variant, and even at this concentration, no signal was detected at the array (Figure 3B). To assess whether this behavior could be reverted by enforced tetramerization of the receptor, we also generated and tested the double mutant GR^P474R/P625A^. Variant GR^P474R^, also known as GR^tetra^, forms tetramers both in the nucleoplasm and at the array (Figures 3A-B) (24). Although introduction of this exchange in the DBD did not modify the monomeric character of the GR^P625A^ mutant in the nucleoplasm, it did rescue tetramerization when bound to DNA, indicating a complex, intertwined relationship between DBD and LBD domains within the FL-GR. Dimerization of mutant GR^Y545E^ was also impaired in the nucleoplasm, although to a lesser extent (ε = 1.54), as well as tetramerization at the array (ε = 3.27) (Figure 3B).

To verify whether disruption of the non-canonical dimerization interface also affects the biological activity of the receptor, we performed RNA-seq analyses of cells stably expressing variant GR^P625A^. The results of these transcriptomic analyses revealed an almost complete loss of transcriptional activity of the GR^P625A^ mutant (Figure 3C). Furthermore, the nine hypersensitive genes activated by GR^P625A^ showed significantly lower levels of stimulation than the wild-type (WT) receptor (Figure 3D). Altogether, these results indicate that the parallel, Y545-centered GR homodimer is the major active conformation in living cells, and that receptor dimerization is essential for DNA binding and transcription. Because a recently presented structure of multidomain GR features an LBD dimer in which residues of the L6-7 loop and H7 dock onto the N-terminal end of helix H10 (PDB 7PRW (54)), we also generated and studied by N&B the GR^Y640A^ variant. This mutant remained essentially dimeric in the nucleoplasm. Unexpectedly, truncation of the Tyr640 side chain to alanine promoted the formation of higher-order oligomers on DNA (ε = 6.0, indicative of either hexamers or mixtures of higher and lower oligomers; see also below) (Figure 3B). We had previously reported a similar behavior for the Chrousos syndrome mutation, GR^D641V^ (52).

### The Tyr545-dimer is compatible with current structural information on interdomain contacts

The findings presented above strongly suggest that the L1-3 stretch centered on Tyr545 and stabilized by L5-6 residues (e.g., Pro625), is essential for GR homodimerization *in vivo*. This, along with some features of the 7PRW structure, such as the small area and polar character of the LBD-LBD’ interface, prompted us to re-interpret this structure (Figure 4A; Supplementary Video S1). We note that one of the DBD-LBD pairs in 7PRW features an open but well defined L9-10 loop that together with the N-terminus of H10 leans on residues of the C-terminal DBD helix, Arg479 and Gln483 (Figure 4B). The negative charge of the Glu705 carboxylate is compensated by the guanidinium groups from both the preceding Arg704 and Arg714, together with Arg479 (Figure 4B). In addition, residues Arg460–Arg462 at the tip of an exposed loop in the DBD, which play important roles in inter-DBD interactions, occupy a shallow depression on the LBD surface formed by the aromatic side chains of Trp610, Tyr613 and Tyr660 (Figure 4C).

**Figure 4 Legend.**
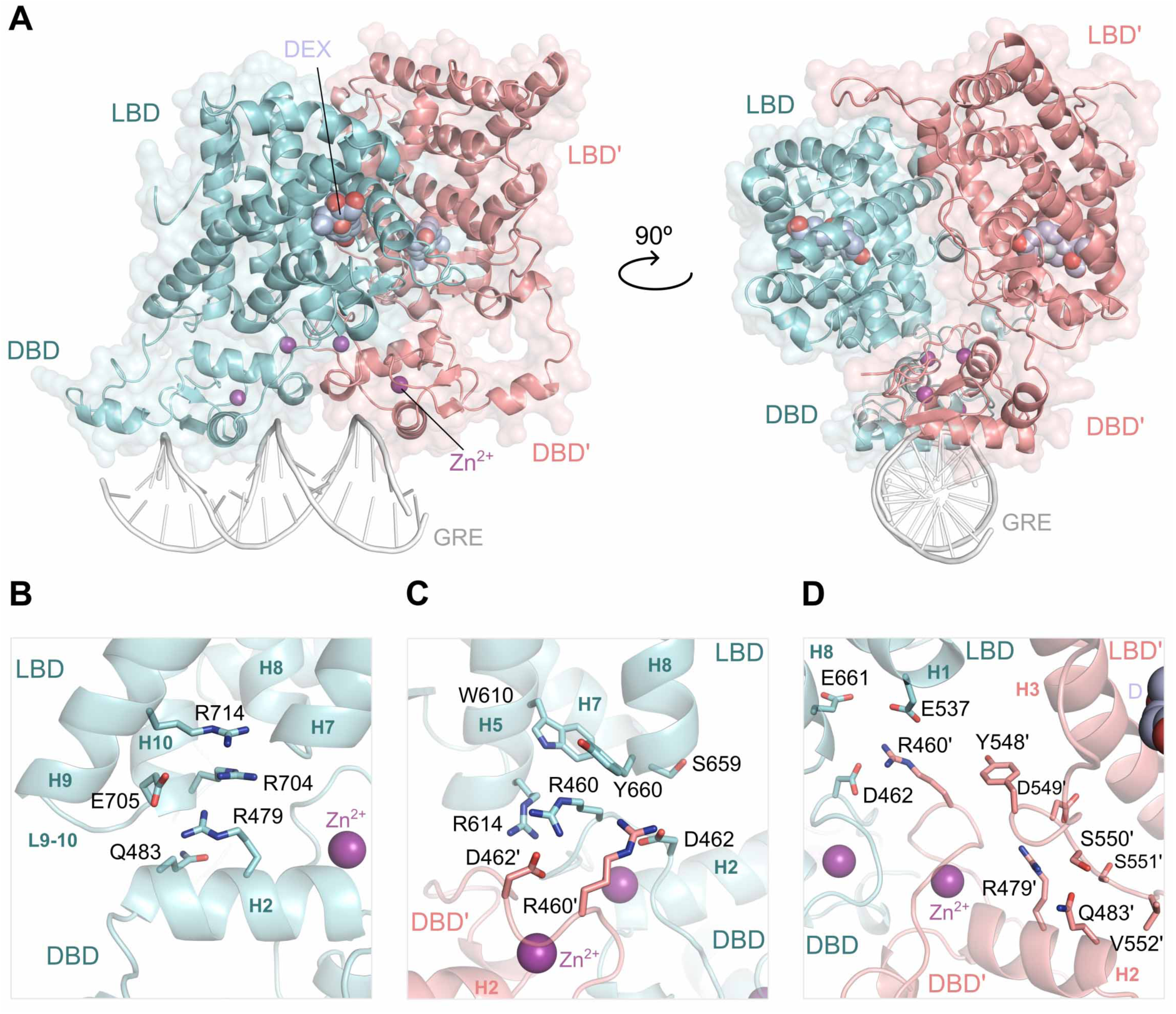
The Tyr545-centered non-canonical dimer of LBD modules is compatible with the canonical DBD homodimer. (A) Two orthogonal views of the 3D model of DNA-bound multidomain human GR generated by superimposing the Tyr545/545’ dimer on the recently reported crystal structure featuring two DBD and two LBD modules (PDB 7PRW (54)). The canonical DBD-DBD’ homodimer and the ‘west’ DBD-LBD pair in 7PRW were left unmodified, while the ‘east’ LBD’ module was repositioned into the non-canonical, Tyr545/545’-centered conformation. Monomers are shown as cartoons colored cyan and salmon, respectively, and DEX molecules are color-coded as in Figure 1. (See also Supplementary Video 1). (B-D) Close-ups of the different interdomain interfaces. (B) At the DBD-LBD interface, the L9-10 loop of the ‘west’ GR monomer is stabilized in the open conformation through contacts with its own DBD. Note that the carboxylate of the conformation-determining residue, Glu705, engages in electrostatic interactions with two neighboring arginines (Arg704 and Arg714) as well as with Arg479 in the DBD module. (C) Contacts between charged LBD and DBD’ residues contribute to dimer stability. Note the projection of Arg460 (DBD’) and Arg614 (LBD) side chains into the neighboring domain. These interactions contribute to stabilizing the DBD-DBD’ interface, especially the symmetric Arg460-Asp462’ / Arg460’-Asp462 salt bridges. (D) Predicted interactions at the DBD’-LBD’ interface. Note that the same residues of the ‘east’ DBD’ module are important for LBD’ binding, most notably the exposed side chains of Arg479’ and Gln483’ (compare to panel B). However, DBD’ would contact residues at the C-terminal end of L1-3 in LBD’ such as Tyr548’ and Asp549’. The Arg460’ guanidinium group is located close to the carboxylates of Glu537 and Glu661, suggesting that additional interactions at the DBD’-LBD interface would contribute to stabilizing the Arg460’-Asp462 salt bridge.

By contrast, the second LBD moiety in 7PRW is located over 15 Å away from the corresponding DBD module. Repositioning of this LBD monomer in the parallel Tyr545/Tyr545’ conformation generates a homodimer in which the more C-terminal stretch of the L1-3’ loop in the second monomer (Tyr548’–Val552’) interacts with the same residues contacted by the ‘west’ GR-LBD module, Arg479’ and Gln483’ (Figure 4D). In this manner, the Tyr545-dimer satisfies the constraints imposed by the canonical, DNA-bound DBD homodimer, while allowing for important DBD’-LBD’ interactions of the ‘east’ GR monomer. Moreover, this model predicts favorable DBD’-LBD interactions as well. In particular, the Arg460’ side chain would compensate for the negative electrostatic potential of Glu537 and Glu661, in addition to the salt bridge with Asp462 in the DBD (Figure 4D). The DBD-LBD/DBD’-LBD’ homodimer interface is conceptually reminiscent of the convergence zone introduced by Rastinejad and co-workers (62).

### Dimer-dimer contacts reveal two alternative mechanisms of oligomeric arrangements with implications for receptor multimerization in living cells

Having established the central role of Tyr545-centered, non-canonical homodimers in living cells, we next explored the possible relevance of observed dimer-dimer interactions for receptor multimerization. We reasoned that the high solvent content of the current crystals results in arrangements that are not imposed by crystal packing contacts and are thus likely to reproduce interactions characteristic of *bona fide* FL-GR multimers detected in living cells, as visualized in our N&B studies (24, 52). Despite their apparent complexity, the arrangements of these GR-LBD dimers in the current crystal can be comprehensibly described by two different combinations of dimer-dimer contacts, which are essentially made by the most flexible regions of the domain, the H6/L6-7/H7 area (Supplementary Figure S4; referred to as the ‘Pro637 setting’) and the H9/L9-10/H10 area (Figure 5, labeled as the ‘Trp712 setting’, see also Figures 2A-B). These two settings not only result in different overall arrangements of the eight GR-LBD dimers in space but also turn out to have radically different functional implications, as discussed below.

**Figure 5 Legend.**
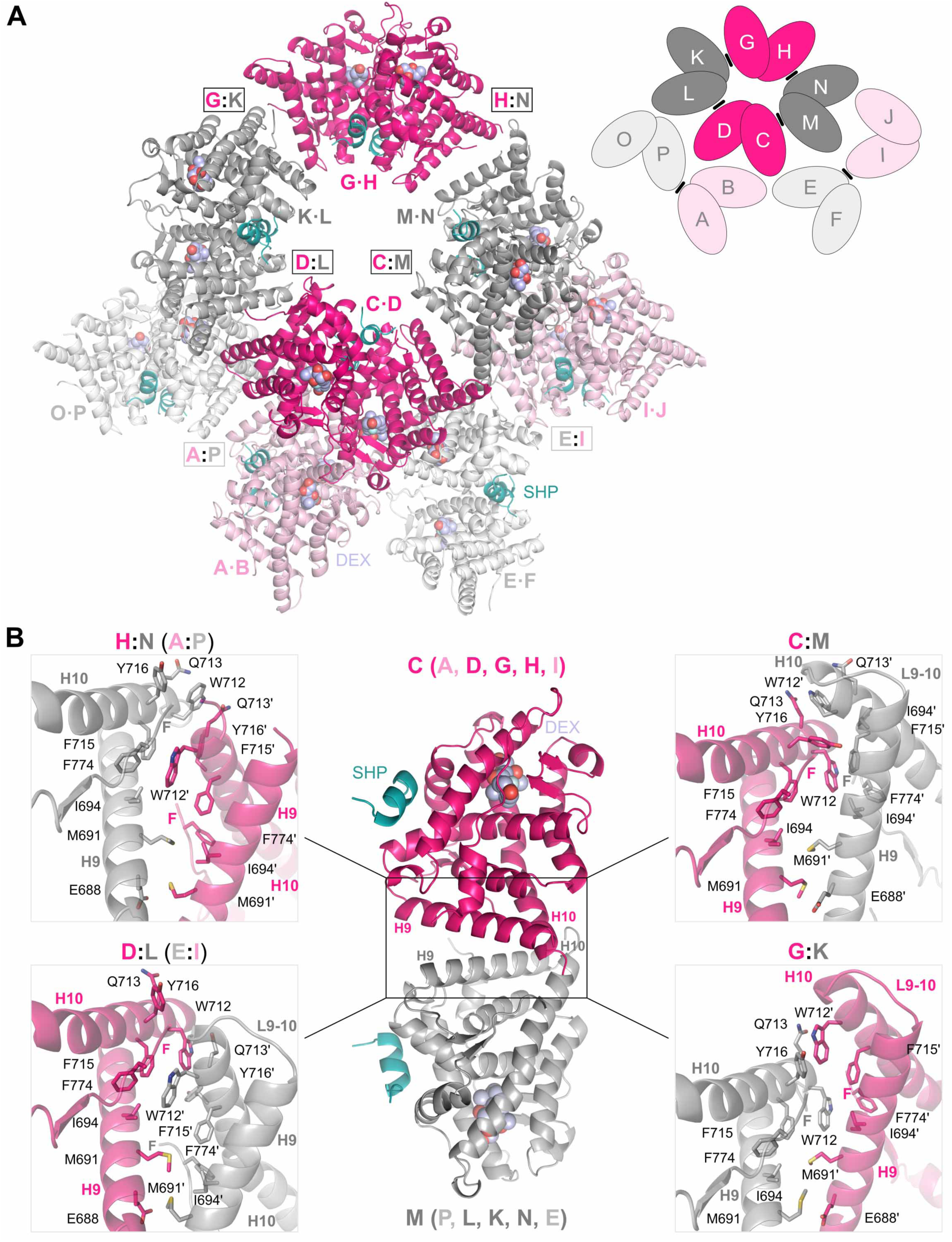
The Trp712 setting features six topologically equivalent dimers-of-dimers generated by contacts between aromatic residues at the N-terminus of H10. (A) Overall view of the eight GR-LBD homodimers in the ASU chosen to highlight dimer-dimer contacts between residues of the H9/L9-10/H10 area, the Trp712 setting. Dimers are colored according to their relative positions in the 3D structure. In the forefront, dimers C·D / G·H and K·L / M·N are colored in fuchsia and dark gray, respectively. In the background, the second layer of dimers, A·B / I·J and E·F / O·P, are colored in pale pink and light gray, respectively. The six inter-dimer contacts centered on the triplet of aromatic residues, Trp712 / Phe715 / Tyr716 (A:P, C:M, D:L, E:I, G:K, and H:N) are boxed. A schematic representation of the setting is shown in the inset to the right, with GR-LBD molecules given as ovals and contacts at the Trp712 interfaces highlighted with black bars. The SHP coregulator peptides and DEX moleculas are color-coded as in Figure 1. (B) Comparison between the six different arrangements of Trp712-centered dimers-of-dimers. The D·C:M·N dimer-of-dimers is shown in the central panel. (For simplicity, only the contacting monomers, C and M, are represented). Close-ups of the six interfaces are shown in the insets: A:P / H:N and E:I / D:L (to the left) as well as C:M and G:K (to the right). Major interface residues are shown as color-coded sticks and labeled. Note the central position of residues Trp712 / Trp712’ and the different inter-monomer orientations (see also Supplementary Video 3).

### A complex network of protein-protein interactions centered on the Pro637 interface generates octameric GR-LBD assemblies

The choice of the ASU that results in a more compact packing of monomers in the crystal focuses on contacts mediated by residues of the L6-7 loop (Leu636, Pro637) and the N-terminus of H7 (e.g., Tyr640, Asp641), with additional contributions from neighboring L1-3/H3 and L11-12 residues (Supplementary Figure S4A and Supplementary Video S2). These contacts, described in detail in the legend to Supplementary Figure S4, generate two similar, irregular octamers, comprising monomers (A·B):(C·D):(E·F):(G·H) and (I·J):(K·L):(M·N):(O·P), with central dimers B:C and J:N, respectively. These tetramers-of-dimers can be considered to form by the initial association of two Tyr545-centered dimers, A:B and C:D, through symmetric contacts of their L6-7 loops (Supplementary Figure S4B and Supplementary Video S2). This is followed by the docking of two additional dimers, E·F and G·H, from ‘below’ and ‘above’ onto this core dimer-of-dimers, respectively (in the orientation shown in Supplementary Figure S4B). Although monomers E and H also rely on L1-3 and L6-7 for protein-protein interactions (PPIs), these ‘lower’ and ‘upper’ subunits dock asymmetrically onto the central dimer-of-dimers so that they simultaneously contact both monomers B and C. The octamer is closed by minor contacts between the peripheral monomers, A-F and D-G. Altogether, 2,165 – 2,200 Å^2^ are protected from bulk solvent at the inter-monomer interfaces. Similar contacts are made by the topologically equivalent GR-LBD modules that form the second octamer, although there are large differences both at the level of relative intermonomer orientations and details of PPIs (Supplementary Figure S4). These observations underscore the high degree of plasticity of inter-monomer contacts in the current crystal. Although this view of the ASU is crystallographically more appealing, it must be stressed that several residues of the Pro637 interface are atypical for biologically relevant protein-protein interfaces, most notably Pro637 itself.

### The Trp712 interface is a plastic, highly populated protein-protein interaction surface

Six of the protein-protein interfaces in the current crystal structure involve residues of helix H9, the N-terminus of H10, and the connecting L9-10 loop, making it the second most common interface (Figure 5A). The most relevant element of this interface is a strictly conserved triplet of nearby aromatic residues, Trp712, Phe715, and Tyr716, arranged around the indole rings of Trp712/Trp712’ (Supplementary Video S3). For this reason, in the following we will refer to this PPI surface as the ‘Trp712 interface’. A comparison of the six Trp712 interfaces reveals common structural features. In the orientation shown in Figure 5B, the indole ring of Trp712 from the ‘left’ GR-LBD molecule docks onto the Gln713’ side chain from the ‘right’ monomer making important VdW contacts. The corresponding Trp712’ side chain from this right subunit projects into an aromatic/aliphatic pocket shaped by the Ile694 (H9) and Phe715 (H10) side chains, along with the phenol ring of the C-terminal Phe774 (F domain). This core interface is completed by additional VdW contacts between side chains of other residues in the central region of H9/H9’, most notably Met691/Met691’ (e.g., Met691-Glu688’, Ile694-Met691’). Altogether, the H9/L9-10/H10 interfaces bury surface areas (BSA) of similar magnitude as *bona fide*, physiologically relevant PPI interfaces(63–65) (470–750 Å^2^) and have large calculated binding energies (Supplementary Table S3).

Although the six Trp712 interfaces are topologically equivalent, as they are formed by the same secondary structure elements, large differences in the inter-dimer orientations are evident (Figures 5B and supplementary S5, and supplementary Video S3). Closer inspection reveals that two of the pairs (A:P/H:N and D:L/E:I) are most closely related to each other, with an overall root mean square deviation (RMSD) of about 2 Å for all Cα atoms (Supplementary Table S4). This leaves four easily distinguishable conformations of Trp712-centered dimers-of-dimers, exemplified by pairs H:N, D:L, C:M and G:K (Figure 5B). Even in two pairs of dimers that adopt similar relative orientations, A:P and H:N, the side chain of Tyr716’ swings between a Trp712-distant and a Trp712-close conformation (Supplementary Figure S5A). Thus, the interconversion between the different Trp712-centered assemblies of GR-LBD tetramers can be described as a ‘screwing’/’unscrewing’ of two GR-LBD dimers relative to each other around the fixed pair of central Trp712 side chains, with additional side chains adjustments (Supplementary Figure S5B and supplementary Video S3). For instance, conversion from the A:P/H:N to the G:K pose involves a large rotation of helix H9’ towards Trp712, allowing for extra interactions of its indole ring with Met691’ (Supplementary Figure S5C), while the Trp712’ and Tyr716 side chains engage in additional contacts (Supplementary Figure S5D).

Although not exactly superimposable on any of the current Trp712-mediated dimers, topologically equivalent arrangements of GR-LBD monomers can be identified in several previously reported crystal structures, including the first structures of the module, also bound to DEX (PDB entries 1M2Z and 1P93 (4, 53)) (Supplementary Figure S5E). This is in addition to GR-LBD complexes with the natural hormone, cortisol (4P6X (66)) or with unrelated nonsteroidal GCs (3CLD, 3E7C and 3K22 (67–69)). Altogether, these observations underscore the high degree of plasticity at the Trp712 interface.

One important feature of these dimer-of-dimers interfaces is that the gathering of the six exposed aromatic side chains generates pockets that could be occupied by aromatic/aliphatic small molecules. In fact, extra electron densities observed in the current structure could be safely interpreted as polyethylene glycol (PEG) molecules of different lengths derived from the crystallization solution (Supplementary Figure S1G). This adds to previous observations by us and others of small molecules bound at this solvent-exposed surface, which is contiguous to, but topologically different from the BF-3 pocket. How this GR ‘sensor site’ regulates GR biological activity will be discussed elsewhere (Jimenez-Panizo *et al*., manuscript in preparation).

### The Trp712 interface is important for receptor tetramerization on chromatin and transcriptional activity

The identification of several Trp712-centered dimers-of-dimers in the current structure, the realization that these contacts are common in previously reported structures and are also populated in solution (52), and the fact that residues such as tryptophans and tyrosines are enriched in *bona fide* PPI surfaces (63–65) prompted us to assess the possible impact of this arrangement in living cells. Firstly, we performed docking experiments to generate 12,000 different dimers-of-dimers using a state-of-the-art protein-protein docking procedure without imposing any constraints (see Supplementary Figure S6A and Materials and Methods section). These solutions were ranked according to their pseudo-energy values, as previously described (52, 70). The results of these experiments revealed that the Pro637-and Trp712-centered areas are hotspots for homotypic interactions, as indicated by their high normalized interface propensity (NIP) values (Supplementary Figures S6B-C). Overall, we found 296 and 422 docking solutions that are close to the symmetric arrangements centered on Pro637 and Trp712 residues observed in the current crystal, respectively, among them one of the top 20 solutions each (Supplementary Figures S6D-E). Of note, many of the docking solutions are compatible with the experimentally observed dimers-of-dimers.

Next, we generated both single (GR^W712E^) and double mutants (GR^W712S/Y716S^) of the most exposed aromatic residues at the interface. We also generated and tested a variant that truncates the two aromatic residues at the alternative LBD-LBD’ interface, GR^Y640A/Y716S^ (Figure 6A). Similar to the mutants of the Tyr545 interface described above, these variants were correctly expressed and trafficked to the nucleus upon DEX treatment (Supplementary Figure S3A). Unexpectedly, all mutants showed reduced receptor dimerization in the nucleoplasm (ε values ∼1.5, Figure 3B), suggesting an impact on non-canonical, Tyr545-mediated homodimer formation either because of impaired DBD-LBD contacts (Figure 4B) and/or through allosteric modification of the L1-3 loop (see above). Mutant GR^W712E^ formed tetramers at the array (ε = 3.76), while simultaneous replacement of the aromatic Trp712 and Tyr716 residues by the less bulky serine resulted in severely impaired tetramerization (ε = 2.56) (Figure 6B), demonstrating a role of this interface in tetramer formation within the context of the FL-GR. Finally, the GR^Y640A/Y716S^ mutant showed an intermediate behavior (ε = 3.0), thus reverting the tendency to the formation of higher-order oligomers of the GR^Y640A^ variant (compare Figures 3B and 6B).

**Figure 6 Legend.**
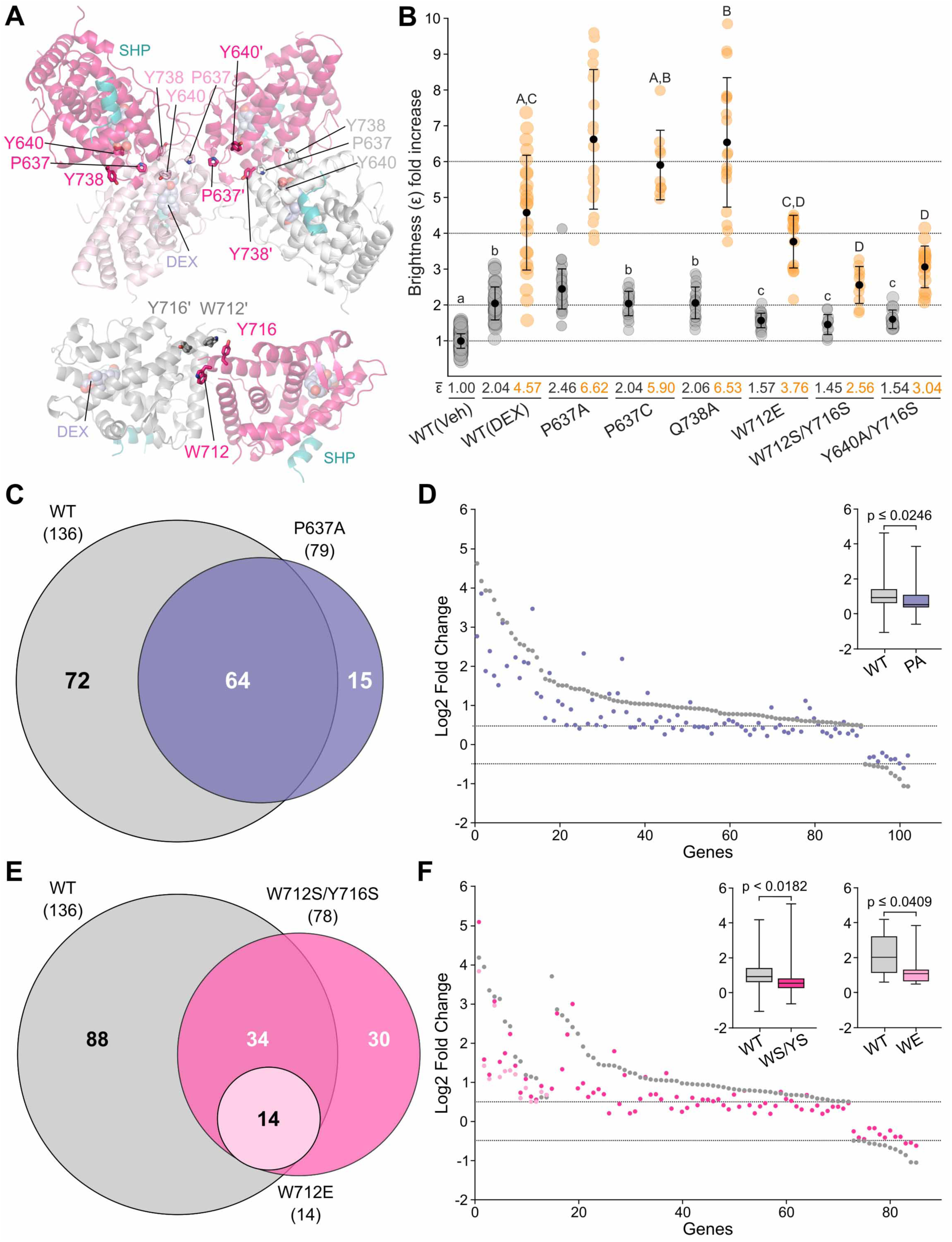
Mutations at Pro637-and Trp712-interfaces differently affect GR multimerization behavior and transcriptional activity. (A) 3D representations of major dimer-of-dimers interactions in the current crystal. The central B/C/E/H tetramer in the Pro637 setting (top panel) and the Trp712-centered dimer (C:M, lower panel) are represented. Interface residues mutated in the current investigation are shown as color-coded sticks and labeled. Residues correspond to the crystallized ancGR2 construct. The SHP coregulator peptides and DEX molecules are color-coded as in Figure 1. (B) Impact of mutations that affect Pro637-and Trp712-interfaces on FL-GR oligomeric state in living cells. The results of N&B assays showing the aggregation state of WT GR and mutants in the nucleoplasm and at the array are represented with gray-and orange-colored dots, respectively. The sample mean is shown as a black point in the middle of each dot plot, together with the standard deviation bars. Dot plots with different superscript letters are significantly different from each other (p < 0.05, one-way ANOVA followed by Tukey multiple comparisons test). Statistical analysis of the nucleoplasm and array data (letters) were done separately (small and capital letters, respectively). (C-F) Transcriptomic analysis of the impact of mutations affecting Pro637-(C, D) or Trp712-interfaces (E, F). Stable cell lines expressing GFP-tagged mouse FL-GR or shown variants were treated with 100 nM DEX for 2 h for RNA-seq analyses. Three replicates were used per condition. Protein-coding genes were considered to be regulated by the GC if they showed an absolute log_2_ fold change (FC) > 0.5 at a FDR < 0.01. Statistical analyses were performed using a two-tailed non-parametric t-test. The total number of DEX-responsive genes is given in parentheses in Venn diagrams (C, E). Dashed horizontal lines in scatter plots (D, F) denote Log2 FC +/-0.5. Box and whiskers plot of the same data displays interquartile range depicting the 25^th^, 50^th^, and 75^th^ percentile as box with the median as black bar. (C, D) Venn diagram and scatter plot comparing genes regulated upon DEX treatment by variant GR^P637A^, as compared to the WT receptor. The scatter plot of shared DEX-regulated genes shows the overall lower transcriptional activity of this mutant. (E, F) Venn diagram and scatter plot comparing genes regulated upon DEX treatment by variants GR^W712E^ and GR^W712S/Y716S^, as compared to the WT receptor. Note the severely impaired transcriptional activity of variant GR^W712E^, pointing to an important impact of electrostatic repulsion between Glu712/Glu712’ residues at the dimer-of-dimers interface and/or enhanced p23-mediated disassembly of transcription complexes. Less dramatic but also important effects were observed with the double mutant, GR^W712S/Y716S^.

Unexpectedly, the transcriptional profile of the GR^W712E^ variant revealed an almost complete lack of transcriptional activity (Figures 6E-F, about 10% of activated genes compared to the WT receptor). On the other hand, the GR^W712S/Y716S^ double mutant was also transcriptionally impaired, although to a lesser extent (Figures 6E-F, ∼60% of the genes). These results illustrate the complexity of the GR structure-function relationship and suggest that the dimers-of-dimers centered on the Trp712 / Phe715 / Tyr716 aromatic triplet play an important role in receptor tetramerization, resulting in gene transcription tempering in living cells (see Discussion).

### Allosteric pathways linking L1-3, L6-7, and L9-10 loops

We have recently shown that a mutation associated with Chrousos syndrome, p.Asp641Val, results in enhanced multimerization of FL-GR in living cells upon DNA binding (52). Indeed, the GR^D641V^ variant formed higher oligomeric structures at the array in N&B assays (ε = 7.19), which can be interpreted as a complex mixture of different stoichiometries including tetra-, hexamers, and octamers. Precisely, octamers formed in the current crystals feature one GR-LBD molecule with an Asp641 carboxylate that docks into an aliphatic pocket formed by the side chains Leu636’/Pro637’ from a neighboring monomer (Supplementary Figure S4B). A valine side chain would engage in more favorable VdW interactions with the apolar Leu636/Pro637 side chains, which provides a straightforward explanation for the increased tendency to form higher-order oligomers by the GR^D641V^ variant. A similar multimerization behavior in living cells was observed for the previously studied variant, GR^P637A^ (ε = 6.62, Figure 6B) (52), as well as for two mutants of the “back” LBD face generated in the current investigation, GR^P637C^ and GR^Y640A^ (Figures 3B and 6B). Therefore, we decided to assess whether other mutations of nearby residues at the back surface also generated higher-order multimers at the array in live cells. Indeed, this was the case for the alanine variant of a more distant residue, Gln738 (ε = 6.53; Figure 6B). These N&B results confirmed that the mutation of residues at the Pro637 interface does not affect GR dimerization, but invariably promotes the formation of higher-order oligomers at the array. Furthermore, transcriptomic analysis of the GR^P637A^ mutant indicated a two-fold reduction in the number of activated genes (Figures 6C-D), similar to our previous results with the Chrousos variant, GR^D641V^ (52).

To explore the possible relevance of other known *NR3C1* point mutations and single nucleotide polymorphism (SNPs) for the structure and function of the GR, we mapped all GR-LBD variants reported to date on the 3D structure of the domain (Figure 7A; major clinical and structure-function information on these variants is summarized in Supplementary Table S2). Noteworthy, several SNPs in addition to Asp641 replace a polar or charged residue by one with an aliphatic or aromatic side chain, not only in L6-7/H7 (e.g., p.His645Tyr) but also in the nearby L11-12 loop (p.Asp742Val, p.Lys743Met, p.Ser746Ile) or the F domain (p.Ser765Leu). This is in addition to another similar mutation linked to PGGR, p.Thr556Ile (Figure 7A) (39, 71). These variants are likely to enhance receptor multimerization, similar to GR^D641V^. Other missense variants cluster in and around well-characterized functional sites, the LBP (p.Met560Arg, p.Ala605Thr) and the AF-2 cleft (e.g., p.Val575Gly, p.Gln597Pro/His/Glu, p.Glu755Val), but also at the dimerization (e.g., p.Ser551Tyr, p.Ile559Asn), and tetramerization interfaces (SNPs affecting positions 689, 697, 699-701 and 703 in H9, 708-711 in L9-10 / N-terminus of H10, and mutants p.Arg714Gln/Leu/Trp in this helix) (Figure 7A). Furthermore, paths cross-connecting the two flexible poles of the module are enriched in disease-linked residues: from L9-10/H10 (Arg714) to the L1-3 loop (Tyr545) and H3 (Arg569) through L7-8 and H5 (the ‘lower path’ in the left panel of Figure 7A), or through H9/H10 and the LBP (the upper path). This is in line with functional studies indicating reduced affinity for DEX and impaired interactions with coregulators for several mutations of residues distant from both LBP and AF-2 (Table S2 and refs. (37, 38, 72–76)). These circuits cross-connecting the major functional areas of GR-LBD are evolutionarily conserved from fish to humans.

**Figure 7 Legend.**
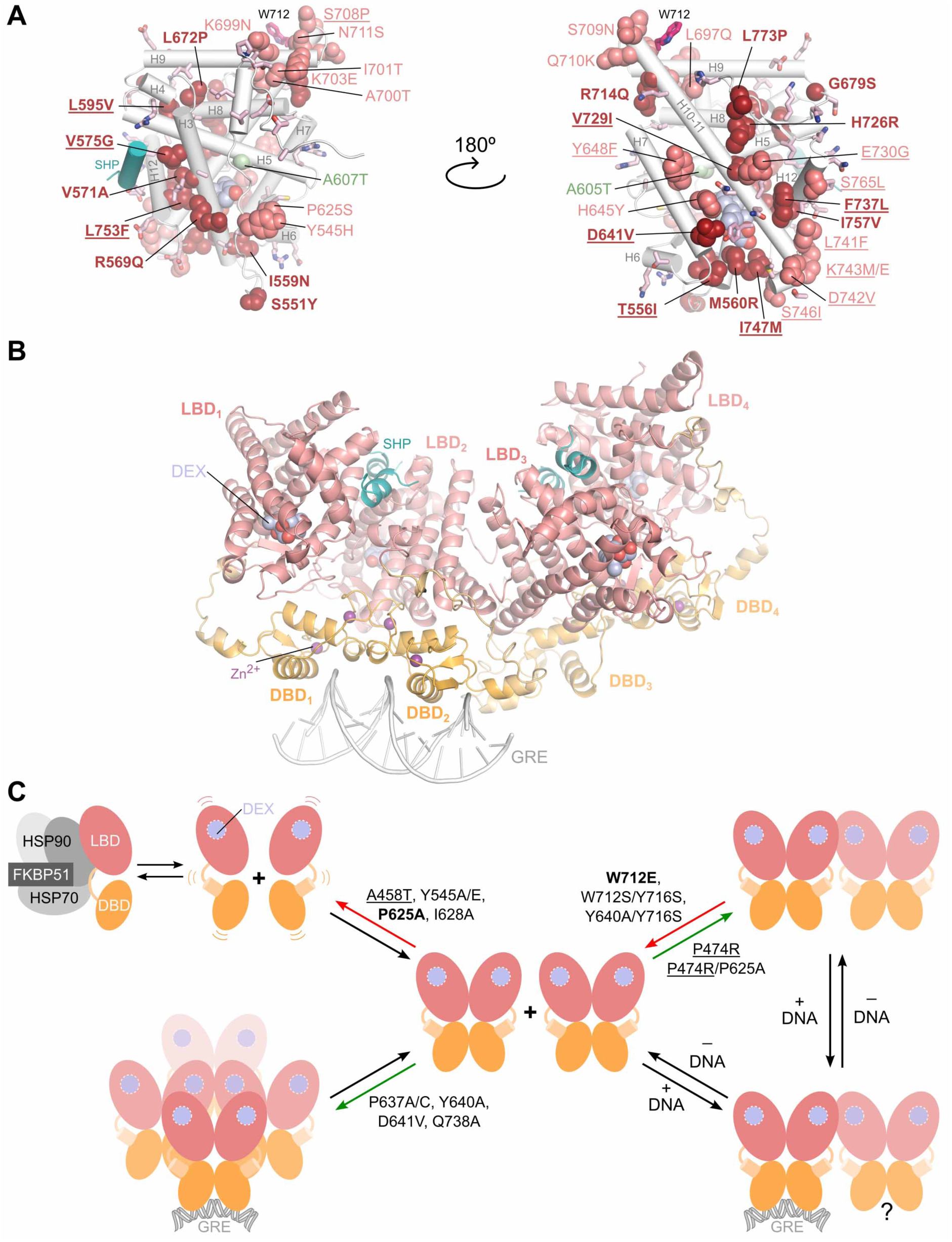
Molecular basis of Chrousos syndrome and suggested pathway of GR multimerization. (A) Distribution of missense mutations and SNPs in human GR-LBD. Front and back faces of the domain are shown in the left and right panels, respectively, with the side chains of affected residues represented as spheres and labeled. Residues have been colored firebrick red to highlight variants identified in patients with Chrousos syndrome, salmon for SNPs that have not been characterized to date but are likely to have an impact on the protein structure and function (e.g., p.Tyr545His, p.Pro625Ser, which are predicted to impair dimerization), and light pink sticks for those SNPs that represent more conservative replacements and are therefore less likely to be deleterious. Variant p.Ala605Thr (green sphere) might increase affinity for GCs, similar to the Ala605Val variant in human (57) and porcine GR (110, 111). Note the clustering of mutations in the functionally relevant L1-3/H3 and H9/L9-10/H10 areas. Note also that several SNPs enhance the hydrophobic nature of the back face (underlined), similar to the previously characterized GR^D641V^ variant (52), and are therefore likely to result in similar oligomerization behavior. (B) Model of tetrameric multidomain GR bound to DNA. All six Trp712-interfaces are compatible with the formation of dimers-of-dimers similar to the one shown in this panel, which corresponds to the A·B:O·P arrangement. Note that tetramerization essentially depends upon LBD-mediated interactions but does not involve contacts between DBD dimers. (C) Proposed pathway leading to GR multimerization and activation of transcription. GR folding requires the coordinated actions of chaperones Hsp70 and 90, cochaperone p23 and immunophilins. Two approaching monomers can dimerize by sequential DBD-DBD’ (through residues of the second zinc finger) and LBD-LBD’ interactions (through the Tyr545 surface). Dimerization stabilizes intra-monomer DBD-LBD and DBD’-LBD’ contacts mediated by residues of the L1-3 or L9-10 loops, respectively, along with additional intermolecular interactions. GR dimers, rather than monomers, can then independently ‘scan’ the chromatin landscape (100), followed by the recruitment of coregulators, which ultimately activates transcription. DNA binding triggers a conformational change, leading to tetramerization through contacts between the Trp712-centered surfaces, which might enhance the transcriptional output. Based on our results, the impact of point mutations on the oligomeric state of the receptor is indicated (DBD mutations are <u>underlined</u>). Mutations that have the highest effect on GR transcriptional activity are in **bold**. For simplicity, association events are presented sequentially, but are likely highly coupled *in vivo*, and therefore most mutations have pleiotropic effects on the multimerization process. Most notably, mutations at the H9/L9-10/H10 zone affect monomer conformation (through impaired contacts with the DBD), dimerization (via allosteric effects transmitted to L1-3), and tetramerization (by impairing dimer-of-dimers formation). Finally, the kinetics of the multimerization process is influenced not only by the concentrations of GR and GCs, but also by the presence of specific coregulators and the chromatin state (not illustrated in the figure for simplicity). See Supplementary Figure 7 for a thorough classification of studied mutants according to their impact on GR oligomerization.

## Discussion

From a biochemical point of view, protein crystals are highly concentrated protein solutions. This feature complicates the biological interpretation of crystal contacts, as these solvent-excluded surface areas are similar to those typical of transient protein-protein complexes. Although the enrichment in specific residues such as Trp, Tyr and Arg as well as geometric considerations allow for tentative assignments of some crystal contacts to *bona fide* PPI interfaces *in vivo* (63–65), further complementary orthogonal evidence is needed to establish the actual biological relevance of these interfaces. On the other hand, analysis of single-point mutants in living cells might be obscured by differences in expression levels and/or in the interaction with chaperones and thus the fraction of well-folded molecules. Furthermore, a given mutation might impact not only the local protein structure but also distant areas of the receptor through allosteric effects. Besides, point mutations can have manifold direct and indirect implications on the function of the studied protein, which for NRs include both inter-domain and inter-monomer interactions, as well as contacts with DNA and with coregulators (77–79). Analysis might be further complicated by the differential impact of mutations on PTMs, as we have recently shown for AR (80, 81).

The current investigation benefits from the unusually high solvent content of grown GR-LBD crystals, which implies that the observed interactions are less conditioned by crystal packing effects than in the dozens of previously reported crystal structures of GR-LBD (52, 82, 83). This is demonstrated by the high degree of plasticity observed at Pro637 and Trp712 interfaces (Figures 5, supplementary S4 and S5; supplementary Videos S2 and S3). Furthermore, the study of a battery of structure-guided point mutants by fluorescence microscopy in living cells and analysis of their impact on GR transcriptional activity allows us to refine the catalog of multimeric GR arrangements and their functional relevance *in vivo* (Figures 3 and 6). Altogether, our current results reconcile a large body of structural and functional information on the receptor gathered over the last two decades. Specifically, it provides conclusive evidence that the L1-3 loop, and in particular the Tyr545 side chain (phenolic ring), correctly positioned to engage in interactions with a neighboring GR molecule by residues of the underlying H5-H6 linker, Pro625/Ile628, is essential for receptor homodimerization in living cells and transcriptional activity. Furthermore, they strongly suggest that the parallel orientation of GR-LBD modules (#6 in our catalog) dominates *in vivo*. Any perturbation of the Ile628/Pro625/Tyr545 cluster (Figure 2; Supplementary Video S1) would report to the dimer interface in a domino-like effect. This explains the relatively minor effect of the Ile628Ala mutant in previous N&B experiments (24), by contrast to the large impact of the central Pro625 residue demonstrated here (Figure 3) and by others (4, 61). Other topologically unrelated arrangements of GR-LBD modules in which two Tyr545 side chains stack on each other, in particular another parallel dimer (#11, see Figure 1 and ref. (52)), are in principle compatible with the canonical DBD homodimer, but would lead to much weaker DBD’-LBD’ contacts. Nevertheless, this conformation might play a role during receptor dimerization. Our results also point to a strong structural and thermodynamic coupling not only between protein-protein dimerization interfaces (DBD-DBD’ and LBD-LBD’), but also with protein-DNA interactions. Thus, the double mutant GR^P474/P625A^ remained monomeric in the nucleoplasm but formed tetramers at the array (Figure 3). In this regard, GR-DBD monomers adopt different relative orientations in free and DNA-bound homodimers (49–51).

Furthermore, inter-dimer contacts revealed in the current structure appear to be relevant for the process of receptor multimerization. Our N&B results indicate that the second most populated interface in the crystals, formed by residues of the H9/L9-10/H10 area, plays a critical role in FL-GR tetramerization (Figure 6). This is also in line with our previous crosslinking studies coupled to mass spectrometry, which identified “top-to-top” dimers that cross-connect residues of H9, L9-10, and the F domain as one of the hotspots of inter-monomer interactions (52). The three nearby, exposed aromatic residues at the N-terminus of H10, Trp712, Phe715 and Tyr716, emerge as a major element of this dimer-of-dimers interface. Probably because of the high degree of flexibility at this interface, we have observed important differences in the impact of point mutations of these aromatic residues. Thus, the previously characterized mutant, GR^W712A^, had only a minor impact on receptor multimerization (52). This contrasts with the large impact on multimerization but in particular on transcriptional activity of variant GR^W712E^, which only stimulated the same set of hypersensitive genes as GR^P625A^, although trafficking normally to the nucleus. This apparent contradiction points to a more profound impact of electrostatic repulsion at this interface, which perhaps reduces the dwell time of the GR^W712E^ variant. In addition, transcription complexes organized around GR^W712E^ would be more efficiently disassembled by p23 and re-assemble less efficiently (84), because the Glu712 carboxylate could form strong, solvent-protected salt bridges with Arg122 in the co-chaperone (11). The observation of a more compact, closed conformation with enlarged helix H10 in these dimers-of-dimers is in line with the common trend of increased secondary structure formation at the expense of irregular structures/loops in PPI interfaces (64, 65). We note that H9 features the sole characterized PTM of the GR-LBD, SUMOylation of Lys703 (85, 86). Modification of this lysine residue was shown to increase GR transcriptional activity (86).

We also note that in several previously reported crystal structures the Trp712 / Phe715 / Tyr716 aromatic triplet engages in important interactions with either the L1-3 loop (homodimers #7 and #8 in our catalog), L6-7 and surrounding residues in H11 (dimer #13) or the C-terminal F domain together with either H9 (homodimer #20) or H11 (dimers #18 and #19). This is in addition to other symmetric arrangements that were not included in our initial classification because of their low BSAs (52). These assemblies feature two topologically distinct, ‘collapsed’ sensor sites (i.e., arrangements with closely packed triplets of aromatic residues): PDB entry 5UC3, corresponding to a dominant negative mutant of GRα (GR^L733K/N734P^) complexed with mifepristone, and 6NWK/6NWL, featuring ancGR2-LBD bound to either DEX or hydrocortisone (87). These diverse arrangements appear to reflect the intrinsic tendency of solvent-exposed aromatic residues of the L1-3, H7, but in particular at the N-terminus of H10 to engage in PPIs and are less relevant for the function of the GR *in vivo*. However, the role of some of these arrangements in certain pathophysiological settings cannot be completely ruled out at present. We also note that the Trp712 interface has been postulated by others as the major GR dimerization interface (88). Our current results suggest that this protein surface is critical for receptor tetramerization, although N&B data implicates an indirect effect in receptor dimerization likely through the allosteric pathways connecting L1-3 and L9-10 loops. This is also in line with our previous N&B results showing dimerization of the (A458T, I628A) double mutant (GR^mon^) at the array (24).

A third pathophysiologically relevant arrangement that emerges from the current results and previous research is centered on residues of the L6-7 loop and the N-terminus of H7, along with nearby residues of L1-3 and H11/L11-12. Alanine truncation of Pro637 (L6-7), Tyr640 (H7) or Gln738 (H11), or replacing the charged Asp641 by an aliphatic valine led in all cases to enhanced receptor multimerization at the array in living cells, with FL-GR forming mixtures of tetra-, hexa-and octamers (Figure 6). Of note, in addition to p.Asp641Val several PGGR-linked mutants and SNPs would increase the apolar nature of the GR-LBD back surface (Figure 7A). It is also noteworthy that the overall surface areas protected at these L6-7-centered interfaces are significantly larger than those of the Trp712 interfaces. Early studies of another L6-7 residue, Cys638, in rat and mouse GR have provided evidence of the role of this loop in GC binding and transcription (24, 60). Because the L6-7 loop is an important element of the hormone entrance channel to the LBP (89, 90), the impact of these mutants on GC binding adds to the deleterious formation of non-productive multimers. Exchanges that increase the hydrophobic character of this surface would trap the receptor in higher order multimers. Given the exacerbated propensity of mutants of the back GR-LBD face for multimerization, however, it cannot be excluded that L6-7-centered complexes are relevant for condensate formation, which has been shown to modulate coactivator recruitment (30, 91). Alternatively, specific sets of dimers-, trimers-and tetramers-of-dimers that utilize different L6-7 combinations might play an independent role in long-range chromatin interactions and chromatin opening (92–95).

Other oxosteroid receptors are known or predicted to rely on topologically equivalent elements for multimerization. Two peripheral AR monomers dock onto the BF-3 regions of the active AR homodimer through their L6-7 loops, with important interactions involving aromatic residues Tyr632, His635 and Tyr640 (numbered according to the equivalent human GR residues) (10, 96). The resulting tetrameric arrangements might explain our previous observation of FL-AR multimers both in the nucleoplasm and at the array (24). On the other hand, the Trp712/Phe715/Tyr716 aromatic triplet is strictly conserved in GR’s closest relative, the MR, and is involved in important contacts in several crystal structures of MR-LBD. However, non-conservative replacements of nearby residues, most notably the Gly698>Arg exchange, favor a completely different, antiparallel arrangement of MR-LBD modules in which the Trp712 side chain docks onto the guanidinium group of Arg698. It is conceivable that these specificities of the H9/L9-10/H10 surface play a major role in receptor multimerization (25, 97), and might underlie formation of different GR–MR heterodimers (98). Altogether, the L9-10 loop together with the surrounding termini of H9 and H10 emerges as a major regulator of multimerization in oxosteroid receptors. Given that the DBDs of all steroid receptors are nearly identical, and the overall conservation of residues that shape the hormone binding pocket, it is tempting to speculate that the unique transcription programs activated by these NRs are largely dependent on the formation of distinct multimeric arrangements on chromatin.

In summary, current structure and function information can be integrated into the following hypothetical pathway of GR multimerization and transcription activation (Figures 7B-C and supplementary S7). GR is maintained in a monomeric, partially unfolded state by interactions with the Hsp90-p23 ‘maturation complex’, in which the Ser519-550 stretch (comprising the hinge, H1, and L1-3) adopts a fully extended conformation (11). After nuclear import and eventual receptor dissociation from the chaperone complex, this N-terminal peptide completes the folding of the GR-LBD module. Several simultaneous interactions prime the newly folded GR molecule to adopt dimerization-competent conformations, most notably intra-LBD allosteric pathways cross-connecting the L1-3 and L9-10 loops and intra-monomer interactions of these loops with the DBD domains. Dimer formation involves synergistic DBD-DBD’ and LBD-LBD’ interactions mediated through the C-terminal zinc-finger modules and the L1-3 loops, respectively, supported by additional inter-monomer interactions at the confluence zone. Once formed, these Tyr545-centered homodimers can independently ‘sample’ the chromatin landscape (27, 29, 31). Current evidence suggests that interaction with a single GRE is enough to trigger transcription (19). However, binding of two GR dimers to spatially proximal DNA sequences in the appropriate relative orientation would allow contacts between their H9/L9-10/H10 surfaces to generate more stable transient tetrameric complexes on chromatin that organize the transcription machinery. Furthermore, these complexes might increase the concentration of active dimers close to chromatin, facilitating transfer of the transcriptional machinery to spatially close transcription start sites. The essential role of the DNA in this process is highlighted by our previous observation that DNA binding triggers GR tetramerization in living cells (24, 26) and its function as an allosteric modulator of GR structure shown by others (19).

An important element of the current model is that receptor multimerization does not proceed as a hierarchically organized series of events but as the result of concerted, multidirectional intra-and interdomain, as well as inter-monomer interactions, which are modulated by contacts of the receptor with coregulators and target DNA sequences. This feature explains the extraordinary impact of mutations at the DBD dimerization interface (P474R, GR^tetra^) on receptor tetramerization, while on the other hand exchanges at the tetramerization site (W712E, (W712S, Y716S)) have a large influence on homodimer formation. Future investigations should establish whether the different dimer-of-dimers centered on the Trp712 interfaces correspond to specific DNA sequences, chromatin structures, and/or coregulators, and activate unique transcription programs. Alternatively, these assemblies might represent intermediates in a pathway involving rearrangements of both DNA and transcription factors that ultimately converge in a single stable transcription complex (99).

## Author contributions

G.L.H., P.F.-P. and E.E.-P. designed and supervised the project, obtained financial support, and share overall responsibility.

A.A.-M., I.M.-N. and M.A. produced and characterized recombinant proteins. A.A.-M. and M.A. performed crystallization trials.

A.A.-M. and I.M.-N. created supplemental videos.

A.J.-P. and J.F.-R. performed and interpreted docking experiments and analyzed structures.

A.A.-M., A.J.-P. and T.A.J. generated FL-GR mutants.

A.A.-M., A.J.-P., A.L.L., T.A.J. and D.M.P. performed and analyzed N&B experiments in cells.

A.A.-M. and E.E.-P. collected X-ray diffraction data.

P.F.-P. interpreted X-ray diffraction data, solved and refined the structure.

P.P. contributed tools and provided information on human *NR3C1* point mutations and SNPs.

T.A.J. performed and with A.A.-M, A.J.-P., A.L.L. and D.M.P. analyzed genomics data.

D.M.P., and G.L.H. supervised cell experiments.

D.M.P. supervised A.L.L and obtained financial support P.F.-P. and E.E.-P. drafted the article.

All authors critically reviewed and approved the manuscript.

## Resource availability

Requests for further information or reagents should be directed to and will be fulfilled by the lead contact, Eva Estébanez-Perpiñá

## Materials availability

All unique/stable reagents generated in this study are available from the lead contact with a completed Materials Transfer Agreement.

## Data availability

The atomic coordinates and structure factors have been deposited at the Protein Data Bank (PDB) with accession code 9HDF and are publicly available as of the date of publication. The Gene Expression profiles are deposited in the Gene Expression Omnibus (GEO) database and the code assigned is GSE282871. All data reported in this paper will be shared by the lead contact upon request.

## Declaration of generative AI and AI-assisted technologies

No generative AI or AI-assisted technologies were used at any stage.

## Acknowledgements

We thank Erick Ortlund (Emory University) for providing the ancGR2-LBD expression plasmid. We thank ALBA-Cells synchrotron XALOC team for beamline support.

## Supplementary data

Supplementary Data are available at NAR Online.

## Abbreviations

AF-2: Activation function-2
AR: Androgen receptor
ASU: Asymmetric unit
BF-3: Binding function-3
BSA: Buried surface area
DBD: DNA binding domain
DEX: Dexamethasone
ER: Estrogen receptor
FDR: False-discovery-rate
FBS: Fetal bovine serum
FL: Full-length
GC: Glucocorticoid
PGGR: Primary generalized glucocorticoid resistance
GFP: Green-fluorescent protein
GR: Glucocorticoid receptor
GRE: Glucocorticoid response elements
KO: Knock out
LBD: Ligand-binding domain
LBP: Ligand-binding pocket
MMTV: Mouse mammary tumor virus
MR: Mineralocorticoid receptor
N&B: Number and brightness
NIP: Normalized interface propensity
NR: Nuclear Receptor
NTD: N-terminal domain
PEG: Polyethylene glycol
PDB: Protein Data Bank
PPI: Protein-protein interaction
PR: Progesterone receptor
RMSD: Root-mean-square deviation
RNA-seq: RNA sequencing
SHP: Small heterodimer partner
SNP: Single nucleotide polymorphism
VdW: Van der Waals
WT: Wild type

