## Supplementary figures and tables for "The multimerization pathway of the glucocorticoid receptor"

### Supplementary Figure 1.

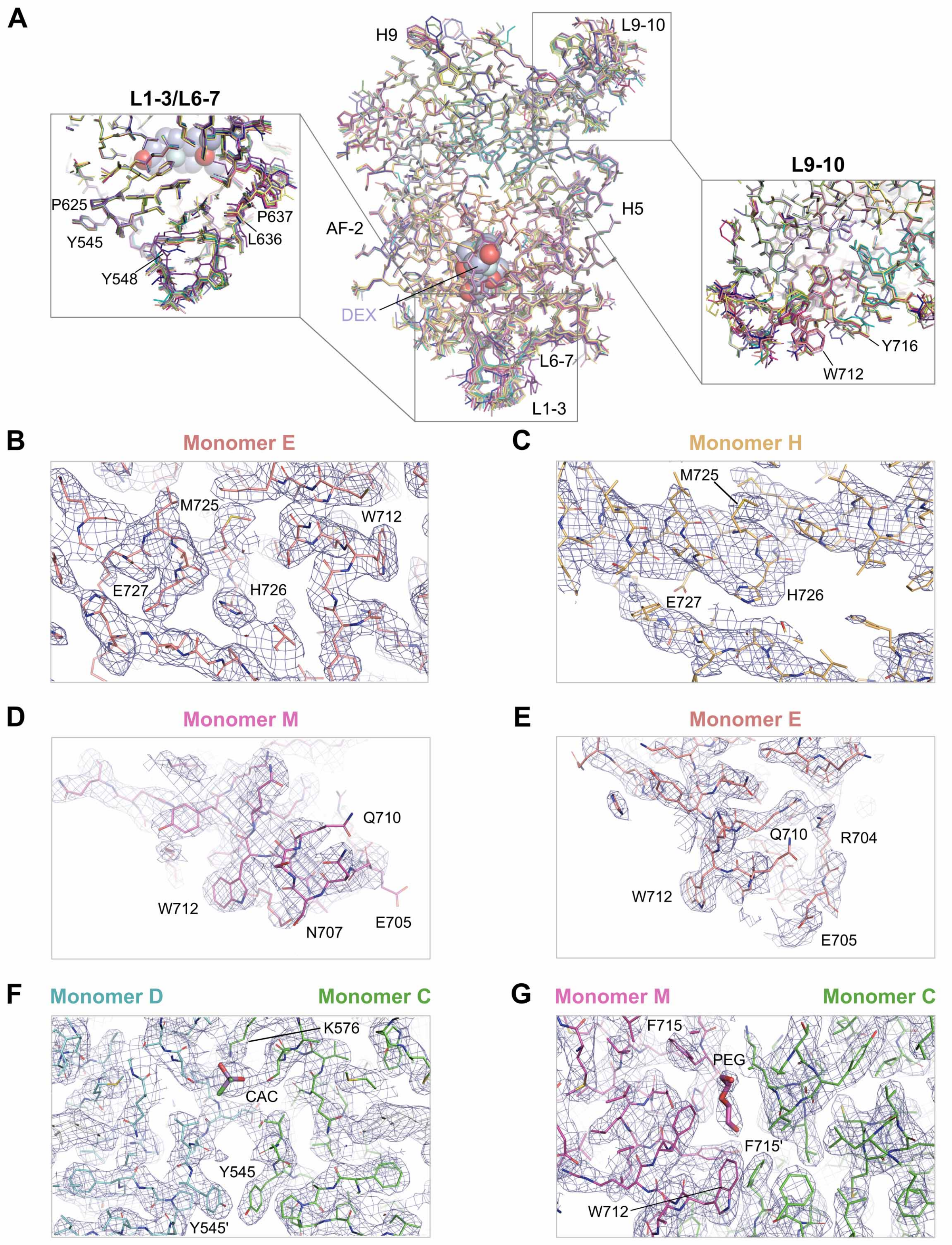

**Supplementary Figure 1 Legend.**

**Details of the current crystal structure, related to Figure 2.**

(A) Superimposition of the 16 crystallographically independent copies of the ancGR2-LBD module found in the ASU of current crystals (center panel). The 16 monomers are denoted A to P, with the SHP peptides that occupy their AF-2 clefts in a canonical conformation termed a to p, respectively. Note that structural variability is limited to two protein regions: (i) the C-terminal end of loop L1-3, together with loops L6-7 / L11-12 and the C-terminal end of helix H11, located at the bottom of the domain in this orientation (left inset), and (ii) loop L9-10 together with the N-terminus of H10, at the opposite pole of the domain (right inset). The latter features a triplet of solvent-exposed, strictly conserved aromatic residues, Trp712, Phe715, and Tyr716.

(B – G) Details of the final model with the $2F_{obs} – F_{calc}$ electron density map superimposed, contoured at 1 σ. Both main- and side chain atoms are shown, and a few residues are labeled.

(B) Close-up around helix H10 of monomer E. Note that both main- and side chain atoms are perfectly defined by electron density. The electron density map is of similar quality in all other monomers, except monomer H.

(C) Detail around helix H10 of monomer H. Note that the side chains of most residues are not defined by electron density, pointing to significantly higher thermal motion. (See also supplementary Figure S2).

(D) Close-up of loop L9-10 and the N-terminus of H10 in the ‘closed’ conformation. In several monomers (exemplified here by monomer M) the side chains of Asn707 and Gln710 stack on each other so that their carboxamide oxygen atoms accept hydrogen-bonds from the main chain nitrogen atoms of the opposite residue. In this manner, H10 is N-terminally extended to residue Asn707 and L9-10 is reduced to a tight turn formed by Glu705 and Gly706, with the acidic Glu705 side chain pointing towards the solvent. Note that the main chain of all residues is well defined by electron density.

(E) Close-up of loop L9-10 and N-terminus of H10 in the ‘open’ conformation. In contrast to the closed conformation shown in panel D, in several monomers Glu705 points towards the H9–H10 interhelical space, which is not compatible with formation of the Asn707 / Gln710 cap. Note that the L9-10 loop is not well defined by electron density in these monomers, indicating enhanced mobility.

(F) Extra electron density at the ‘upper’ interface regions of all basic dimers (A·B, C·D, …, and O·P) could be interpreted as a cacodylate ion (CAC) derived from the crystallization solution, which is trapped between the side chains of residues Ser573/573’ and Lys576/576’.

(G) PEG molecules could be identified occupying several of the inter-monomer pockets created by the triplet of aromatic residues at the N-termini of H10/H10’, Trp712/712’, Phe715/715’ and Tyr716/716’. The molecule bound at the C:M interface is shown here.

### Supplementary Figure 2.

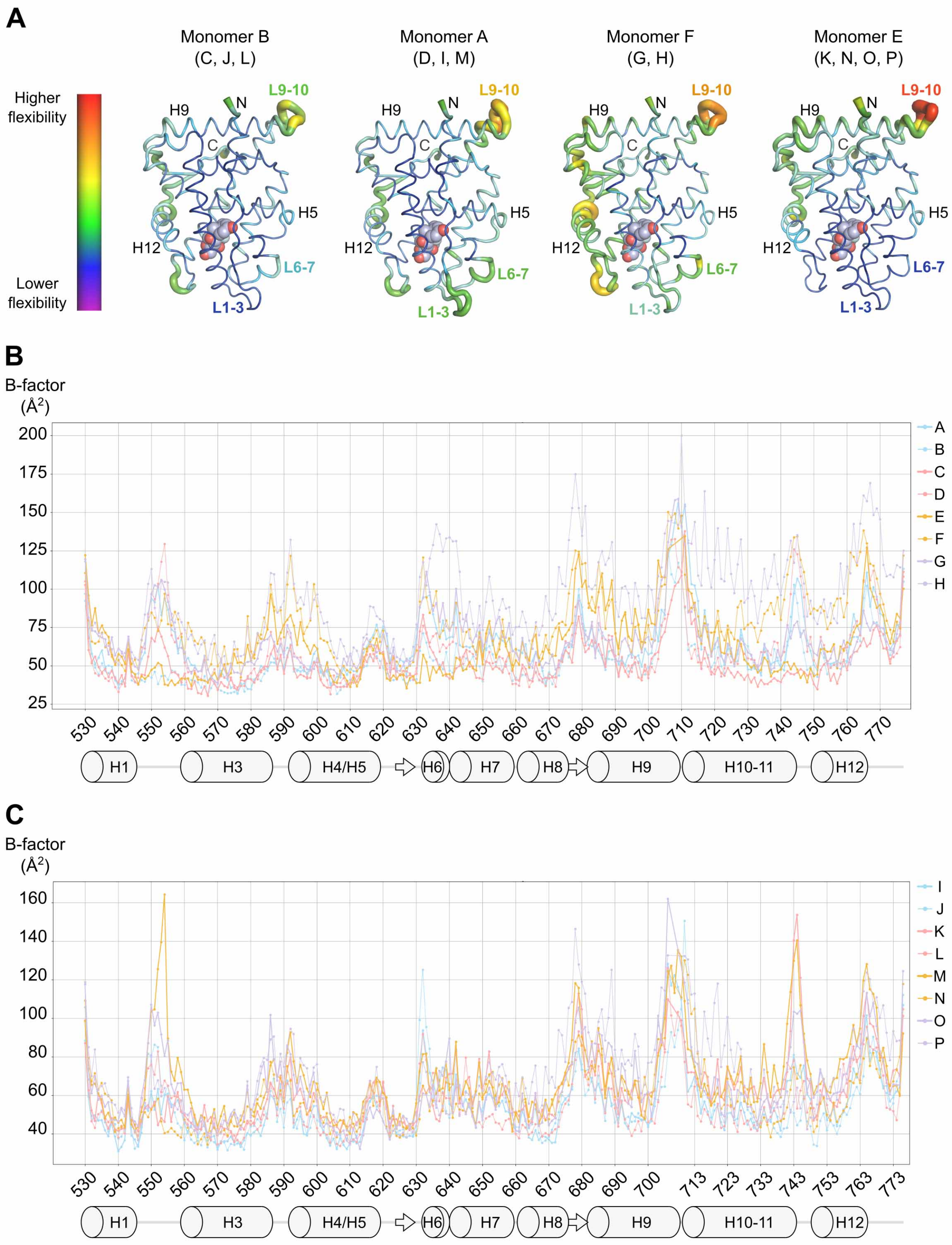

**Supplementary Figure 2 Legend.**

**Intra- and inter-monomer variations in protein mobility, related to Figure 2.**

(A) Qualitative representation of the distribution of temperature factors (B-factors) in the current crystal structure. Four representative GR-LBD monomers are shown as ribbons rainbow-colored and sized according to the average B-factor of each residue (i.e., protein regions characterized by large thermal motions appear thicker and with warm colors (yellow, orange and red), while those with low mobility are thinner and in cold colors (green, blue and violet)). B-factors were set to 200 $Å^2$ for the few residues of the L9-10 loop not defined by electron density in monomers E, H, K, N, O and P. Note that monomers with the highest mobility in L9-10 show the most rigid L1-3 / L6-7 loops.

(B, C) Plot of B-factors for all 16 crystallographically independent monomers. For simplicity, B-factors are plotted separately for each of the octamers found in the ASU in the Pro637 setting. The secondary structure elements are shown in both panels.

(B) Plot of B-factor *vs*. residue number for the octamer formed by GR-LBD monomers A–H.

(C) Plot of B-factors *vs*. residue number for the octamer comprising monomers I–P.

### Supplementary Figure 3.

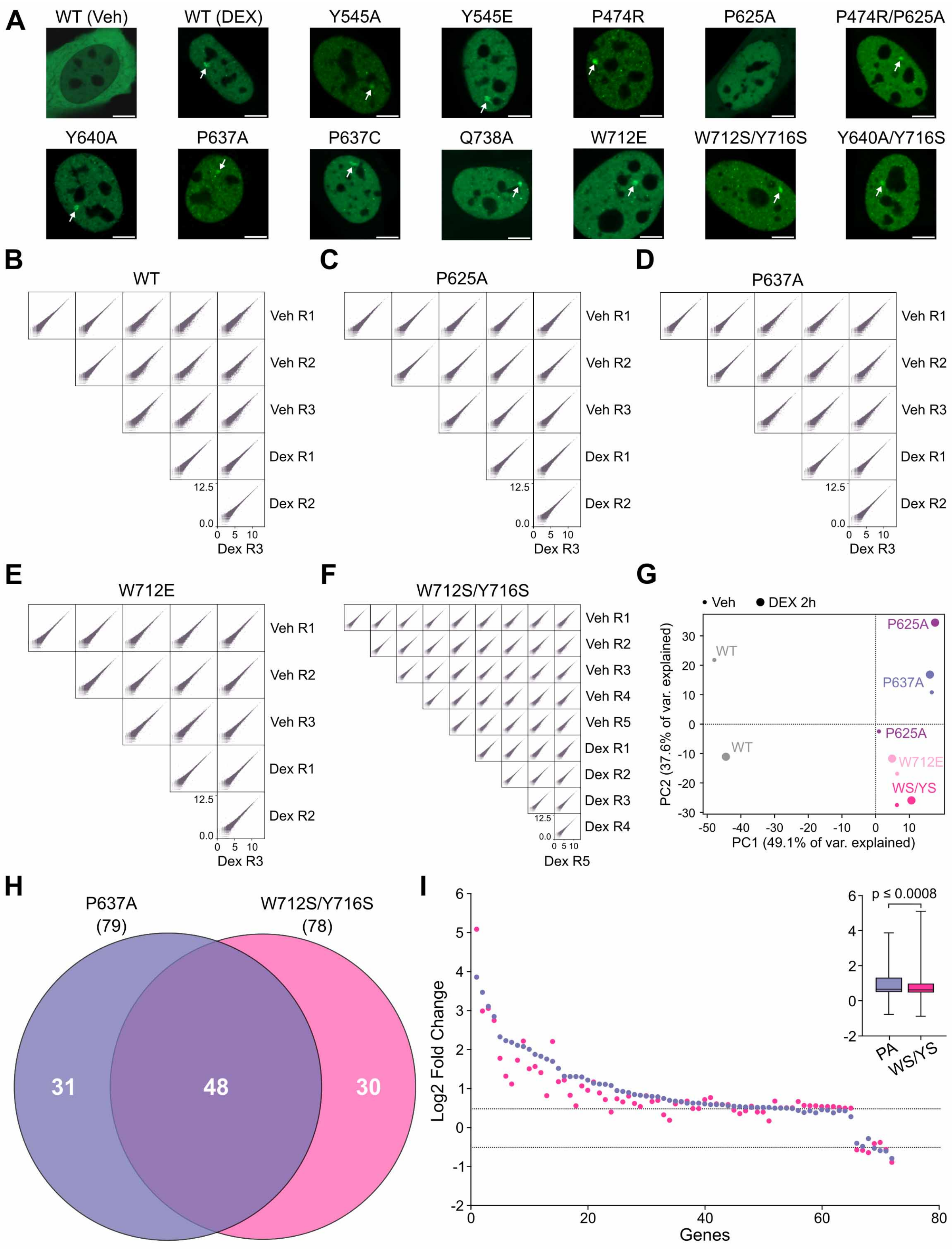

**Supplementary Figure 3 Legend.**

**Quality control of N&B and RNA-seq experiments, related to Figures 3 and 6.**

(A) Subcellular localization of WT GFP-mGR and studied variants in 3617-GRKO cells, as assessed by confocal microscopy. White arrowheads point to the MMTV arrays. Scale bar: 5 µm. N&B experiments have been performed with mouse (m) GR as in previous investigations (see main text for references), but residue numbers correspond to the human receptor to facilitate comparisons.

(B–F) Reproducibility of RNA-seq experiments. Pearson correlation of total RNA-seq replicates (R1-3 or 5) for each GR variant (top label) and treatment (Veh or DEX), as noted (see STAR Methods). All $R^2$ values equal 1.0.

(G) Principal component analysis (PCA) of total RNA-seq data using replicate 1 from each cell line/treatment (see STAR Methods).

(H) Venn diagram comparing hormone-regulated protein-coding genes (2 h DEX treatment/vehicle) in variants $GR^{P637A}$ and $GR^{W712S/Y716S}$. Three replicates were used per condition. The total number of hormone responsive genes (FDR < 0.01, |Log2 FC| > 0.5) is given in parentheses. Note the overall equivalence of hormone-regulated genes in both cases.

(I). Scatter plot of Log2 FC for the responsive genes shared between the two variants. Dashed horizontal lines denote Log2 FC +/- 0.5. Box and whiskers plot of the same data displays interquartile range depicting the $25^{th}$, $50^{th}$ and $75^{th}$ percentile as box with the median as black bar. Statistical analysis was performed using a two-tailed non-parametric t-test.

### Supplementary Figure 4.

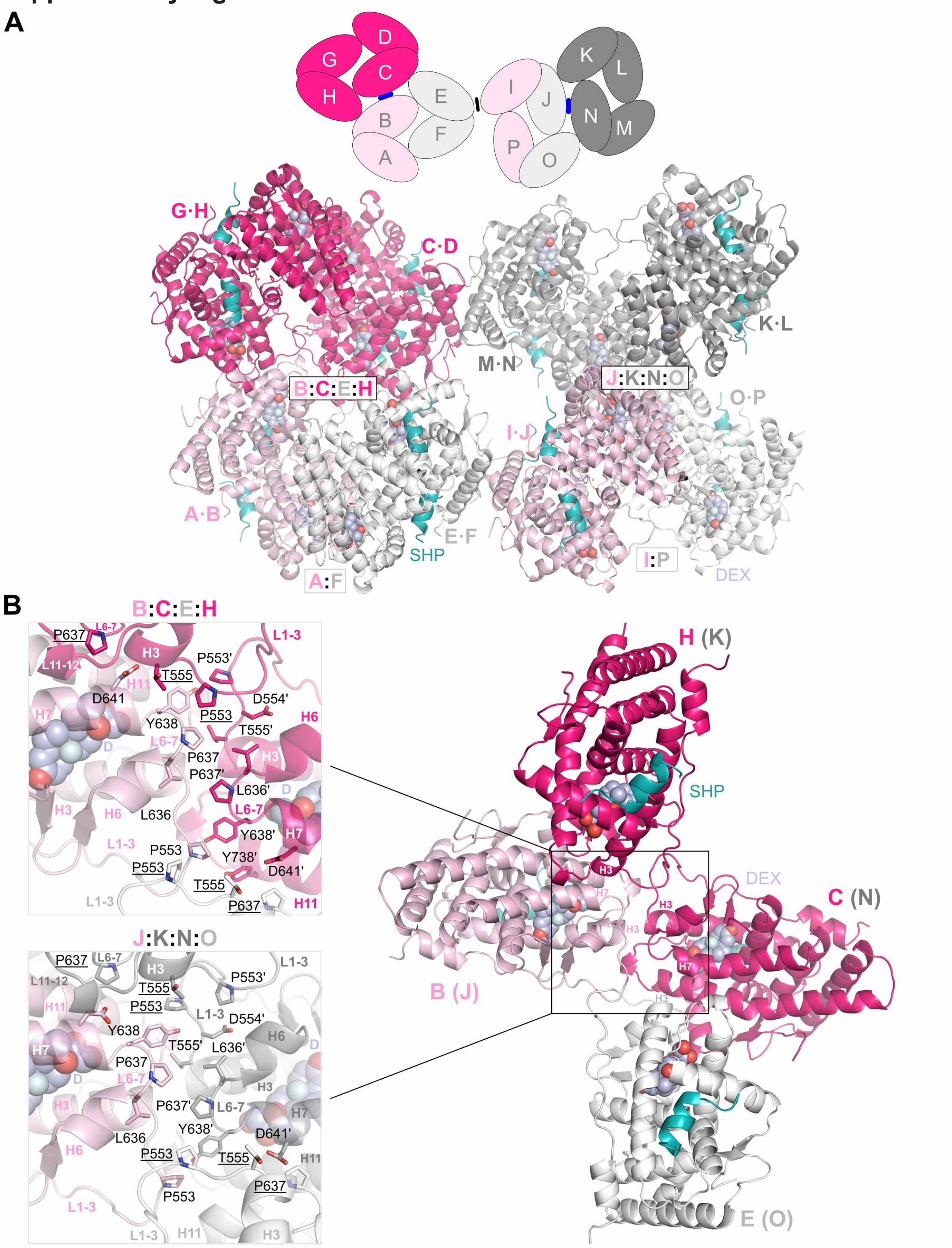

**Supplementary Figure 4 Legend. The Pro637 setting focuses on the interactions in the L6-7/H7 area, related to Figures 2 and 5.**

(A) Overall view of the eight GR-LBD homodimers in the ASU chosen to highlight dimer-dimer contacts between residues of the L6-7/H7 area, the Pro637 setting. Dimers are colored according as in Figure 5. The SHP coregulator peptides and DEX molecules are color-coded as in Figure 1. The central 'convergence zones' dominated by PPIs between the L6-7 loops of monomers B/C/E/H and J/K/N/O are boxed. (Inter-dimer contacts are denoted with colons). A schematic representation of the setting is shown in the inset on top of the panel, with GR-LBD molecules given as ovals and contacts at the Pro637- and Trp712-interfaces as blue and black bars, respectively. (The second Trp712-centered contact, C:M, is lost in this 2D projection). Note that in this setting the ASU is divided into two equivalent octamers, composed of monomers A-H and I-P, respectively. Although these two octamers are topologically equivalent they differ considerably in the intermonomer orientations and the details of PPIs. Most notably, the distances between the closest atoms of the central Leu636–Pro637 pairs increase from < 4 Å in the B:C dimer to > 5 Å at the J:N interface. This is accompanied by large relative displacements of the 'upper' and 'lower' dimers (monomers K and O, respectively), which nevertheless are synchronized to allow the interactions of the peripheral domains, I-P and L-M. (See also supplementary Video S2).

(B) Detail of the central E:(B:C):H tetramer (right panel),. Close-ups to the left show the side chains of residues that form the convergence zones in the two tetramers as color-coded sticks. All residues are labeled and labels of residues from the 'upper' and 'lower' monomers are underlined. Briefly, two Pro637/637' side chains oppose each other and contact the Leu636'/636 aliphatic side chain from the neighboring monomer. These interactions are strengthened by several additional contacts between the N-termini of H3/H3’, which dock onto the L11-12 loops of the neighboring monomer. The Pro637/Leu636 residues of the upper and lower GR-LBD moieties dock onto helices H7 (Asp641, His645) and/or H11 (Tyr735, Gln738) of the central dimer. (Phe735 / Tyr738 in the ancGR2 sequence used for crystallization. For simplicity, we refer to the natural human residues). Additional interactions lock together the tips of the L1-3 loops of pairs B-E and C-H (from Ser550 to Asp554). The total BSAs at these trimer interfaces range between 1,250 and 1,460 $Å^2$. Finally, the peripheral monomers of the upper and lower dimers (i.e., those not involved in L6-7-mediated contacts, A-F and D-G), engage in additional contacts through their C-terminal L1-3 residues. Although the areas protected at these peripheral interfaces are certainly too low to be significant in isolation *in vivo*, they contribute to generating stable octamers and allow for communication between the N-terminal L1-3 loops of the peripheral GR-LBD monomers.

### Supplementary Figure 5.

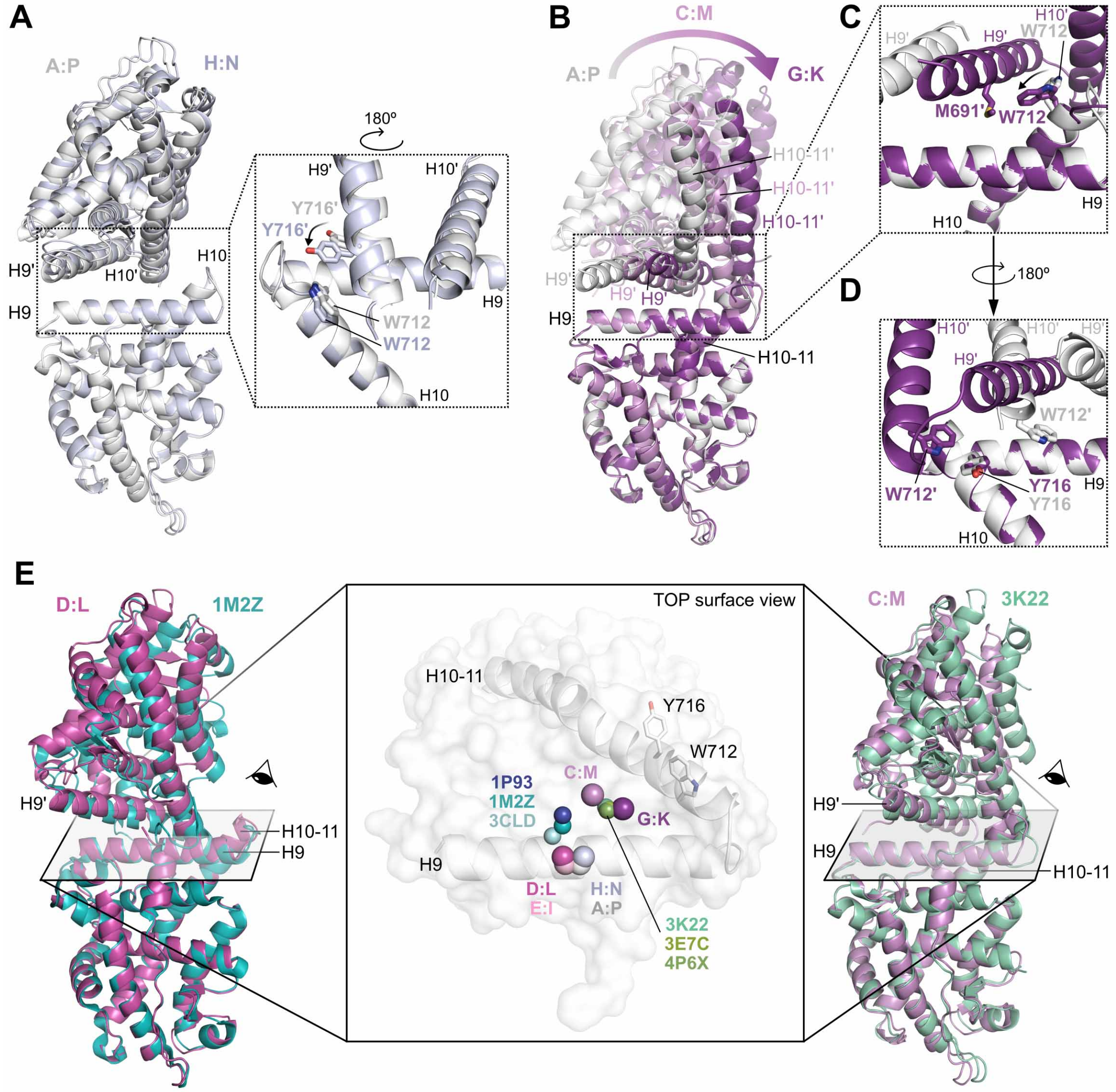

**Supplementary Figure 5 Legend. The Trp712 dimers-of-dimers interface features a high degree of plasticity, related to Figures 2 and 5.**

(A) Superimposition of A:P and H:N dimers, represented as cartoons colored light gray and blue-white, respectively. The central H9/H9’ and H10/H0’ helices are labeled. Note the small but significant displacements of residues in the upper GR-LBD monomers relative to each other, when the lower monomers are used for superimposition. Note also that these differences are not limited to a rotation of the upper monomer as a rigid body, but also include large variations in side chain conformations such as flipping of the Tyr716’ side chain from a Trp712-distant to a Trp712-proximal position (inset).

(B) Trp712-centered dimers-of-dimers sample a large conformational space. All six crystallographically independent GR-LBD dimers are superimposed on their lower monomers. The central H9/H9’ and H10/H10’ helices are labeled. Note the large rotation made by the upper monomers in C:M (violet) and G:K dimers (dark violet), relative to dimer A:P.

(C and D) Close-ups showing the Trp712-centered interface in dimers A:P and G:K viewed from the front (C) and the back (D) of the lower monomers in the orientation shown in panel B. Note the large displacements of helices H9’ and H10-11’ relative to each other.

(E) Comparison with previously reported crystal structures. A top view of the lower monomer is shown as a white cartoon with superimposed semitransparent surface in the central panel. The centers of mass of Trp712-centered dimers are indicated with spheres colored as above for dimers observed in the current structure, and in different blue/green colors for previous work. Two Trp712 dimers are found in PDB entries 1M2Z, 3CLD, 3K22, and 3E7C, while four are present in entries 1P93 and

4P6X. For simplicity, however, only one homodimer from each entry has been included in the analysis. Note the separation of the current Trp712-centered dimers in four clusters (A:P/H:N, C:M, D:L/E:I, and G:K). Note also that previously deposited structures do not exactly match any of the current dimers, highlighting the large plasticity at this interface. Representative examples are compared to the current crystal in the left and right panels: 1M2Z dimers are most closely related to the D:L/E:I arrangements, while the conformation observed in 3K22 crystals is most similar to the G:K dimer.

### Supplementary Figure 6.

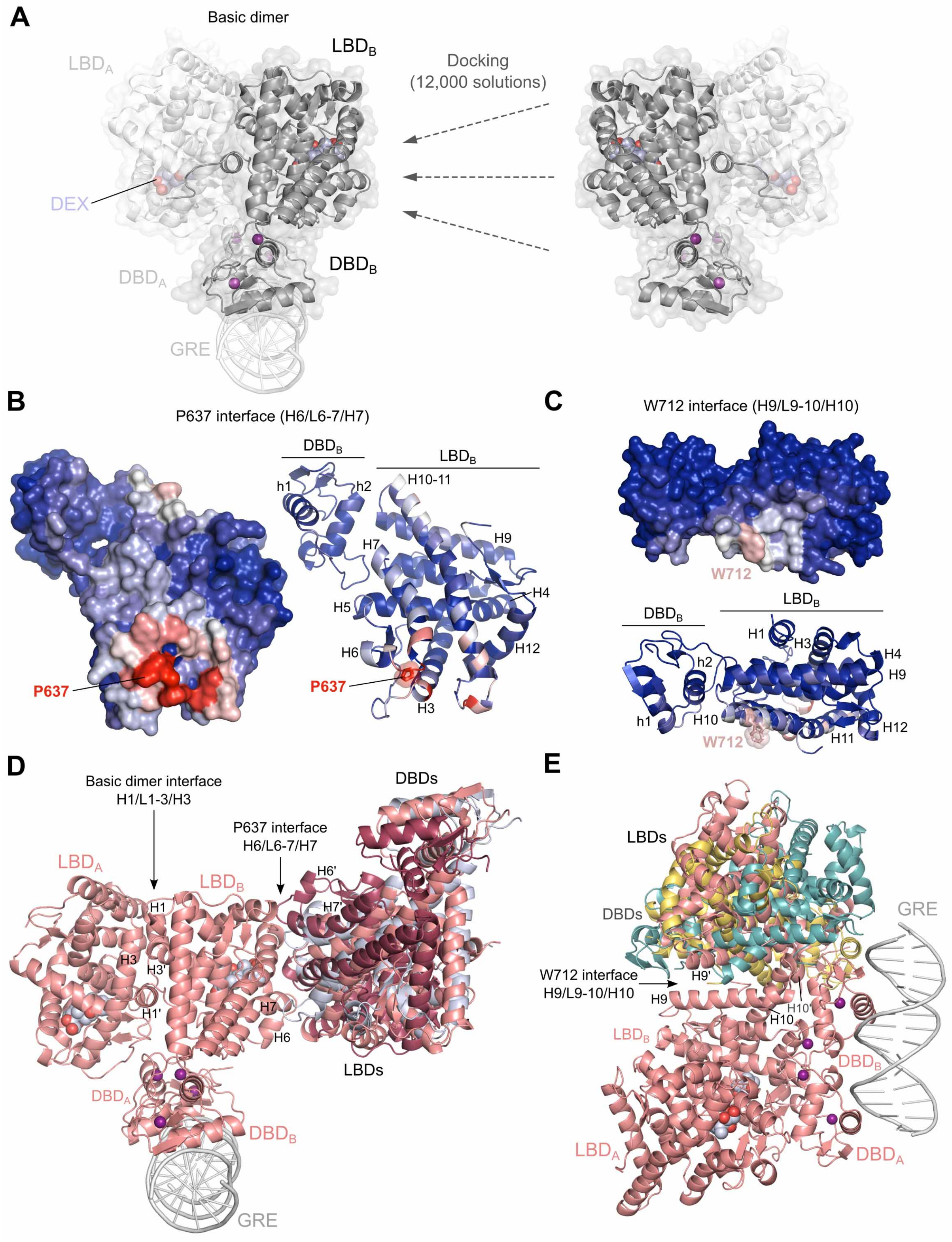

**Supplementary Figure 6 Legend.**

**Docking experiments highlight the propensity of surface areas around L6-7/H7 and L9-10/H10 for homodimerization, related to Figures 6 and 7.**

(A) To reduce bias, a total of 12,000 docking solutions were generated by docking a GR DBD-LBD monomer on the model of the multidomain homodimer described above (see main text and Figure 4) using a state-of-the-art protein-protein docking procedure (see STAR Methods for details).

(B, C) Based on the generated docking solutions, we calculated the normalized interface propensity (NIP) values to identify hotspots of protein-protein interactions. Residues with NIP > 0.4 and < 0 are colored red and blue, respectively, and intermediate values are scaled accordingly. Note that back and top interfaces emerge as hotspots for dimers-of-dimers interfaces that are close to either (B) Pro637- or (C) Trp712-centered assemblies observed in the current crystal structure.

(D, E) Examples of docking solutions close to GR-LBD dimers-of-dimers observed in the current structure. Root-mean-square deviation (RMSD) cutoff: 35 Å. Note that RMSDs for some of the dimers-of-dimers actually found in the structure are about 20 Å (Supplementary Table S4).

(D) A total of 296 docking solutions are centered on residues of the Pro637 interface. In terms of energy, the best solution is ranked top #12 with an RMSD of 14.8 Å, while the one most similar to the Pro637-centered tetramer differs by only 6.6 Å (ranked #1048). The best and the closest docking solutions are shown as raspberry- and lavender-colored cartoons, respectively, superimposed on the model of multidomain GR homodimer.

(E) A total of 422 docking solutions are centered on residues of the Trp712 interface. In terms of energy, the best solution is ranked top #3 (RMSD: 31.7 Å), while the one closest to the Trp712-centered dimers-of-dimers differ by 12.5 Å (ranked #1302). The best and the closest docking solutions are shown as teal- and yellow-colored cartoons, respectively, superimposed on the model of dimeric multidomain GR.

### Supplementary Figure 7.

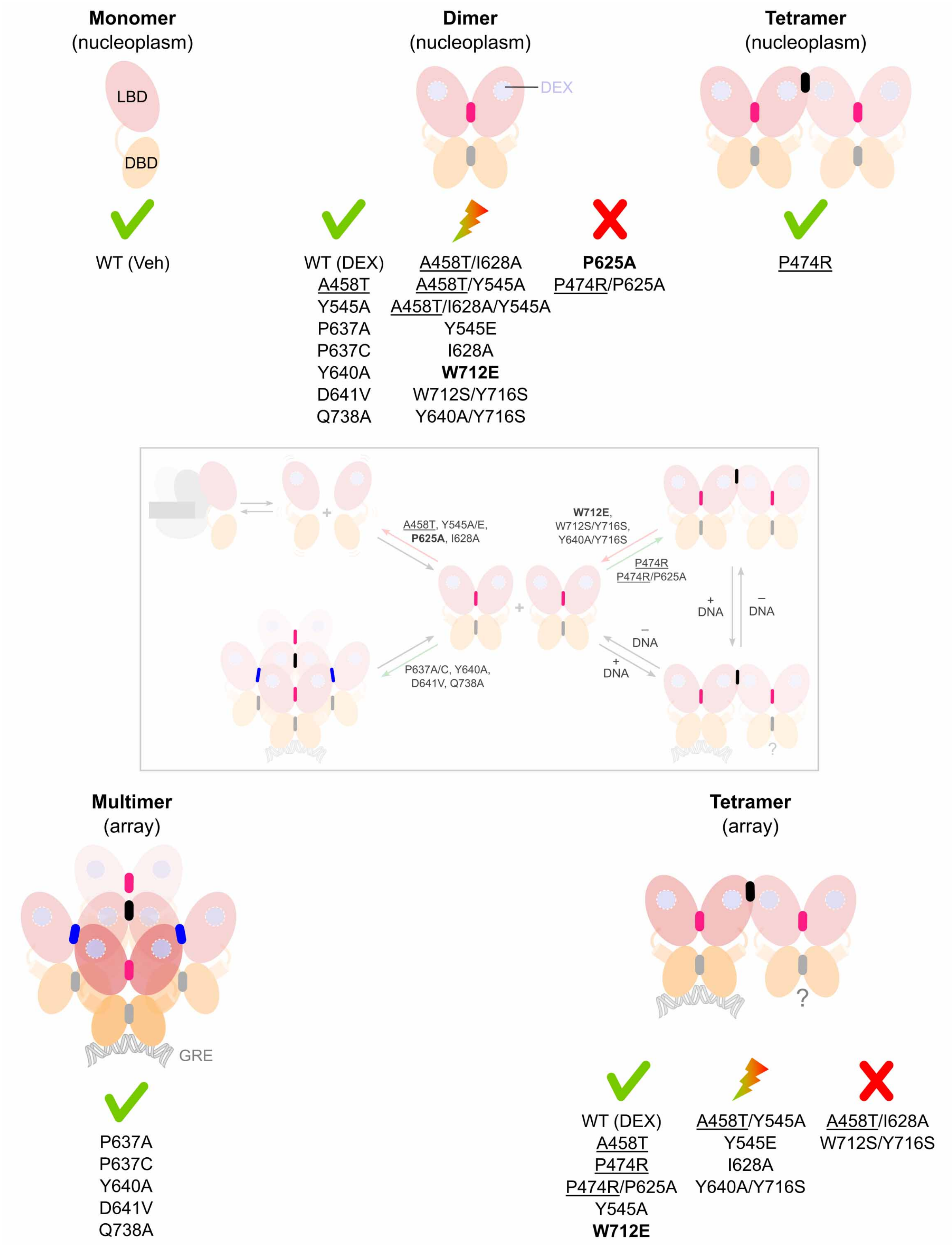

**Supplementary Figure 7 Legend. Impact of studied mutations on GR multimerization, related to Figures 3, 6, and 7.**

The impact of single-, double- and triple-point mutants studied here or in our previous work on the oligomeric state of FL-GR is separately given in the nucleoplasm (upper) and the DNA array (lower panel). GR variants were classified into three categories, highlighted as follows: a check symbol indicates no significant impact, a lightning symbol for mutations having a significant effect compared to DEX-stimulated WT-GR, and a cross symbol for a complete impairment of the corresponding oligomeric form. Mutations are listed below the corresponding group in each case. Interfaces affected are shown as bars colored gray and pink for canonical DBD-DBD' and Tyr545-centered LBD-LBD' interfaces, respectively, black for the Trp712 interface, and blue for Pro637-centered interactions. The proposed pathway leading to GR multimerization and activation of transcription (Figure 7C) is reproduced at the center of the figure for reference. As in Figure 7C, DBD variants are underlined, and mutations that have the highest impact on GR transcriptional outcome are highlighted in **bold**. Neither variant $GR^{P625A}$ nor the triple mutant $GR^{A458T/I628A/Y545A}$ were detected at the array.

**Supplementary Table 1 Legend.**

**Summary of X-ray diffraction data, refinement statistics and model quality.**

| | |
|---|---|
| **PDB code** | 9HDF |
| **Wavelength (Å)** | 0.979 |
| **Resolution range (Å)** | 101.36–2.78 (2.83–2.78) |
| **Space group** | $P2_12_12$ |
| **Cell constants; a, b, c (Å) / α, β, γ (˚)** | 264.41, 265.46, 109.67 / 90, 90, 90 |
| **Molecules in the ASU** | 16 |
| **Total reflections** | 1,115,095 (36,613) |
| **Unique reflections** | 187,084 (8,378) |
| **Multiplicity** | 6.0 (4.4) |
| **Completeness (%)** | 96.8 (88.9) |
| **Mean I/sigma(I)** | 9.9 (1.0) |
| **Wilson B-factor** | 64.998 |
| **R-meas** | 0.140 (1.881) |
| **R-pim** | 0.054 (0.803) |
| **CC1/2** | 0.996 (0.318) |
| **Reflections used in refinement** | 185,038 |
| **Reflections used for R-free** | 1,946 (1.0) |
| **R-work** | 0.21 |
| **R-free** | 0.24 |
| **Number of atoms (protein, DEX, water)** | 34,585 (31,992, 432, 13) |
| **RMS (bonds)** | 0.007 |
| **RMS (angles)** | 1.607 |
| **Ramachandran favored (%)** | 97.06 |
| **Ramachandran allowed (%)** | 2.94 |
| **Clashscore, all atoms** | 7.79 ($99^{th}$ percentile) |
| **MolProbity score** | 2.13 ($98^{th}$ percentile) |
| **Average B-factor** | 68.427 |

**Supplementary Table 2 Legend.**

**Summary of human *NR3C1* point mutations and SNPs affecting the GR-LBD.**

Variants associated with familial or sporadic primary generalized glucocorticoid resistance (PGGR, also known as Chrousos syndrome) are highlighted in bold. SNPs have been taken from https://www.ncbi.nlm.nih.gov/snp. ACTH, adrenocorticotropin hormone; AF, activation function; GC, glucocorticoid; GRE, glucocorticoid-response element; MMTV, mouse mammary tumor virus; NCOA2, nuclear receptor coactivator 2 (formerly GR-interacting protein 1, GRIP1), NCOR1, nuclear receptor corepressor 1. Transactivating activity has been assessed by the MMTV promoter.

| Variant (cDNA[1] / Protein[2]) | Clinical findings / Pathogenicity | Structural and functional impact | Ref. |
| --- | --- | --- | --- |
| c.1586C>T / p.Thr529Ile (SNP rs1818305078) | Not reported in ClinVar. Probably benign. | Replaces polar Thr almost strictly conserved from amphibians to humans by an aliphatic Ile. Position most commonly occupied by Val in ray-finned fishes. Unlikely to have a large impact on the protein structure and function. | – |
| c.1589C>T,G / p.Pro530Leu,Arg (SNP rs756140407) | Not reported in ClinVar. Probably pathogenic. | Replaces well conserved, exposed proline at the N-terminus of the LBD by residues with bulky side chains: Leu -reported in some sequences from cartilaginous fishes- or Arg. Might impair interactions with the maturation complex. Might affect secondary interactions with coregulators. | – |
| c.1591A>G / p.Thr531Ala (SNP rs752542414) | Not reported in ClinVar. Probably benign. | Truncation of a well-exposed threonine strictly conserved from ray-finned fishes to humans. Pro, Ser and exceptionally Ala reported in sequences from cartilaginous fishes. Might have a minor impact on the protein structure and function. | – |
| c.1592C>G / p.Thr531Ser (SNP rs1338961582) | Not reported in ClinVar. Likely benign. | Conservative replacement of a well-exposed threonine strictly conserved from ray-finned fishes to humans by the Ser almost strictly conserved in the MR. Unlikely to have a significant impact on the protein structure and function. | – |
| c.1609G>A / p.Glu537Lys (SNP rs1818299255) | Not reported in ClinVar. Probably benign. | Reversal-of-charge replacement of a well-exposed glutamate strictly conserved from amphibians to humans by the lysine strictly conserved in fishes. Residue almost strictly conserved in the MR. Might slightly impair interactions with the DBD module from a neighboring molecule in Tyr545-centered homodimers. (Repulsion from Arg460'). | – |
| c.1612G>A / p.Val538Ile (SNP rs912945617)[3] | Not reported in ClinVar. Probably benign. | Position 538 is invariably occupied by either Val (from reptiles to humans) or Ala (in ray-finned fishes). Both Ala and Val are found in amphibians, | – |

[1] Numbers refer to mRNA transcript variant 1, NM_000176.3.
[2] Numbers refer to the full-length sequence of human GRαA (UniProt entry P04150).
[3] Not validated.

| | | | |
|---|---|---|---|
| | | Met and Val in cartilaginous fishes. The extra methyl group of mutant Ile538 might slightly affect inter-LBD interactions in Tyr545-centered dimers and/or secondary contacts with coregulators. | |
| c.1633T>C / p.Tyr545His (SNP rs761964114) | Not reported in ClinVar. Probably benign. | Replaces an aromatic residue almost strictly conserved as Tyr/Phe in both GR and MR by a histidine. Might impair homodimer formation, although to a lesser extent than tested mutants, Tyr545Ala/Glu. | [1] and this work. |
| c.1636G>T / p.Ala546Ser (SNP rs776838455) | Not reported in ClinVar. Likely benign. | Replaces an internal alanine by the other residue commonly found at this position, serine. Unlikely to have a significant impact on the protein structure and function. | – |
| **c.1652C>A / p.Ser551Tyr** | Index patient: 14 y/o male homozygous for the mutation had a severe clinical phenotype (fatigue, polyuria, hypokalemia, hypertension, and precocious puberty). Elevated levels of plasma ACTH, cortisol, progesterone, and other steroid hormones. Eight relatives heterozygous for the mutation were phenotypically silent. | Impaired nuclear translocation. Reduced affinity for DEX (2x). Impaired homodimer formation by about 57%. Decreased ability to transactivate GC-responsive genes (3.2x).<br>This position is invariably occupied by either Ser or Thr. Ser/Thr are also most common in the MR, although Ala has also been reported. The bulkier tyrosine side chain might interfere with receptor maturation due to enhanced interactions with the maturation complex. Might interfere with productive DBD-LBD interactions. | [2] |
| c.1666A>G / p.Thr556Ala (SNP rs938836578) | Not reported in ClinVar. Probably benign. | Although Ser/Thr dominate at this position, Ala is also found in several GR sequences and is most common in the MR. Might have a minor impact on the domain structure and function. (Decreased stability of the L6-7 loop). | – |
| **c.1667G>T / p.Thr556Ile** | Patient: 56 y/o male heterozygous for the mutation with a right adrenal incidentaloma and PGGR. Significantly elevated serum cortisol levels, which could not be suppressed by low dosage of DEX. Normal blood pressure. | Delayed nuclear translocation (3.4x). Reduced affinity for DEX (2x). Preserved ability to bind to DNA and to interact with NCOA2 (mostly through AF1). Reduced ability to transactivate GC-responsive genes (2x), but 30% increase in the ability to trans-repress NF-κB-target genes. No dominant negative effect upon GRα-mediated transcriptional activity. Likely to result in a decreased stability of the L6-7 loop (see above). (The bulkier Ile side chain would clash with residues of the L6-7 loop, most notably Pro637). | [3,4] |
| **c.1676T>A / p.Ile559Asn (SNP rs104893909)** | Patient: 33 y/o male carrying a heterozygous mutation with infertility and hypertension, who developed pituitary Cushing's disease at age 38. Causes severe PGGR also in the heterozygous state. | Negligible ligand binding and extremely weak transcriptional activity. The mutant receptor showed a strong dominant-negative effect on the transcriptional ability of the WT receptor *in vitro*. The GFP-GR(I559N) chimera was predominantly localized in the cytoplasm, and only high doses or prolonged GC treatment triggered complete nuclear import (180 vs. 12 min for WT GRα). Inhibited nuclear import of WT GRα, suggesting that its trans-dominant activity is exerted at the level of nuclear translocation.<br>This position is invariably occupied by an | [5–7] |

| | | | |
|---|---|---|---|
| | | aliphatic residue (Ile in amniotes; Leu in amphibians and fishes). A polar, buried Asn559 would compromise LBD folding, in particular the conformation of the L1-3 loop. Might form non-productive heterodimers with WT-GR. | |
| c.1679T>G / p.Met560Arg (SNP rs1818291506) | Not reported in ClinVar, but likely pathogenic. | This position is invariably occupied by an aliphatic residue (Met or Leu; the latter is strictly conserved in MR sequences). A charged Arg side chain might interfere with hormone binding. Might also affect the conformation of the AF-2 cleft. (By similarity to tested mutants, Met560Leu/Thr). | [8] |
| c.1684A>G / p.Thr562Ala (SNP rs1599769111) | Not reported in ClinVar. Likely benign. | Threonine is most common at this position, but the less bulky Ala/Ser are also found in several mammalian sequences. Unlikely to have an important impact on the protein structure and function. | – |
| c.1685C>T / p.Thr562Met (SNP rs761295829) | Not reported in ClinVar. Probably pathogenic. | A residue with a bulkier side chain at this position might clash with Asp549/Ser551 in L1-3. Might indirectly impair homodimer formation. | – |
| **c.1706G>A / p.Arg569Gln (SNP rs1460349075)** | Patient: 45 y/o male heterozygous for the mutation presented with adrenal incidentaloma with mild autonomous cortisol secretion and undetectable ACTH levels. Elevated cortisol and ACTH without clinical symptoms 6 months after surgery, suggesting PGGR. | Reduced affinity for cortisol (2.7x). Impaired nuclear translocation and decreased expression of target genes in response to cortisol, overcome by higher hormone concentrations. Dominant-negative effect on the transcriptional activity at 10 nM cortisol, which was rescued at 100 nM.<br>Replaces the residue strictly conserved from amphibians to humans by the Gln commonly found in ray-finned fishes. Position strictly conserved as Lys in MR sequences. Might impair homodimer formation by decreasing the stability of the L1-3 loop (loss of H-bonds to the carbonyl O atoms of Tyr545 and Ala546) and by eliminating a salt-bridge to Glu542'. | [9] |
| **c.1712T>C[4] / p.Val571Ala (SNP rs104893911)** | Patient: 9 y/o female carrying the homozygous missense mutation presented with pseudohermaphroditism, hypertension and severe hypokalemia. Very high levels of plasma ACTH and high levels of cortisol and other steroid hormones. Her parents and an older sister were heterozygous for the mutation. | Reduced affinity for DEX (6x) and much lower transcriptional activity (10- to 50-fold). Abnormal interaction with NCOA2. Truncates an aliphatic residue strictly conserved from ray-finned fishes to humans. This buried "regional organizer" regulates differential binding of gluco- and mineralocorticoids. Replacement by Met, strictly conserved in MR, resulted in aldosterone binding. Likely to affect domain folding. | [10,11] |
| c.1715T>C / p.Ile572Thr (SNP rs1176598188)[5] | Not reported in ClinVar. Probably pathogenic. | The position is invariably occupied by an aliphatic Ile/Val both in GR and MR. A polar side chain might interfere with homodimer formation. Might have some impact on the binding of coregulators. | – |
| c.1718C>A / p.Ala573Glu | Not reported in ClinVar. Probably benign. | The position is invariably occupied by either Ala or Ser (in amphibians and fishes). A bulkier Glu | – |

[4] Reported as c.1844T>C in the original publication.
[5] Not validated.

| | | | |
|---|---|---|---|
| (SNP rs1239772923) | | side chain could be well accommodated in an extended conformation, and in fact the position is occupied by bulkier Gln/Arg residues in MR. Might slightly interfere with homodimer formation. | |
| **c.1724T>G / p.Val575Gly** | Patient: 70 y/o male carrying missense mutation in heterozygosis presented with bilateral adrenal hyperplasia but was otherwise asymptomatic. His two affected daughters, aged 41 and 47 years, presented mild hirsutism. | Reduced affinity for DEX (2x). Delayed nuclear translocation (2.6x) but retained ability to bind to DNA. Abnormal NCOA2 interaction (through defective AF-2). Reduced transactivation (33%) with enhanced transrepression of the NF-κB pathway (80%). Does not exert a dominant-negative effect.<br>Replacement of a strictly conserved, exposed Val in the AF-2 cleft by a side chain-less glycine would result in severely impaired coregulator binding. | [12] |
| c.1754G>C / p.Arg585Thr<br>(SNP rs749014287) | Not reported in ClinVar, but likely pathogenic. | Replaces an almost strictly conserved, exposed basic residue by a less bulky Thr. The position is occupied by Arg or Lys in the MR. Might weaken interactions with AF-2-bound coregulators. Might also interfere with Tyr545-mediated homodimer formation. | – |
| **c.1783C>G[6] / p.Leu595Val** | Patient: 16 y/o female carrying the missense mutation in heterozygosis[7] presented with acne and hirsutism since menarche, borderline hypertension, and secondary amenorrhea but no signs of hypercortisolism. High levels of both ACTH and cortisol. | No functional characterization of the mutant protein.<br>Replacement of an internal aliphatic residue strictly conserved in both GR and MR by a less bulky Val might decrease the stability of the LBD. | [13] |
| c.1789C>G / p.Gln597Glu<br>(SNP rs769444170) | Not reported in ClinVar. Probably pathogenic. | Isosteric replacement of a strictly conserved, polar glutamine in the AF-2 cleft. Might impair interactions with coregulators. | – |
| c.1790A>C / p.Gln597Pro<br>(SNP rs1289949538) | Not reported in ClinVar, but likely pathogenic. | Non-conservative replacement of a strictly conserved, polar glutamine in the AF-2 cleft. Likely to impair interactions with coregulators. | – |
| c.1791G>C / p.Gln597His<br>(SNP rs1410262431) | Not reported in ClinVar. Probably pathogenic. | Replaces a strictly conserved Gln in the AF-2 cleft by another polar residue. Likely to impair interactions with coregulators. | – |
| c.1813G>A / p.Ala605Thr<br>(SNP rs1300941336) | Not reported in ClinVar. Likely benign. | In addition to Ala, serine is also quite common at this position and is almost invariably found in MR. Thr is conserved in MR from cartilaginous fishes. Might increase domain stability and affinity for hormones. (By similarity to the engineered A605V variant also identified in porcine GR). | [14–16] |
| c.1819G>A / p.Ala607Thr<br>(SNP rs2151548813) | Not reported in ClinVar. Likely benign. | Although Ala is strictly conserved from amphibians to humans, Ser is almost strictly conserved in fish sequences. Ser and the | – |

[6] c.1915C>G in the original publication.
[7] Cannavò et al. report an additional silent mutation in exon 9 (Asn766→Asn), also in heterozygosity.

| | | | |
|---|---|---|---|
| | | isosteric Cys are also most common in the MR. Might increase stability of the LBD by better filling an internal cavity. | |
| c.1840A>G / p.Arg614Gly (SNP rs1817830784) | Not reported in ClinVar, but probably pathogenic. | This position is invariably occupied by a basic Arg/Lys from amphibians to humans, and also in all MR sequences; Glu and Gln predominate in fishes. Radical truncation to a side chain-less glycine might have an important impact on homodimer formation. | – |
| c.1852G>T / p.Ala618Ser (SNP rs1817829879) | Not reported in ClinVar. Likely benign. | Replaces an alanine by a polar Ser found in several species, including primates. Unlikely to have any structural or functional impact. | – |
| c.1861C>A / p.Leu621Met (SNP rs758151443) | Not reported in ClinVar. Probably pathogenic. | Conservative replacement of fully buried, strictly conserved residue, also conserved in other steroid receptors. Might have a small structural impact. Might indirectly affect hormone binding. | – |
| c.1865G>A / p.Cys622Tyr (SNP rs1172094764) | Not reported in ClinVar. Probably benign. | Replaces a partially exposed cysteine by the bulkier tyrosine most common in amphibians, and also strictly conserved in MR. Phe strictly conserved in cartilaginous fishes. Might have a minor impact on the protein structure and function. | – |
| c.1873C>T / p.Pro625Ser (SNP rs1245538911) | Not reported in ClinVar, but likely pathogenic. | Replaces a proline strictly conserved in both GR and MR by a polar Ser. Might impair the conformation of the Tyr545 side chain and thus indirectly receptor dimerization, although to a lesser extent than the P625A variant. | – |
| c.1895A>G / p.Gln632Arg (SNP rs1816993040) | Not reported in ClinVar. Probably benign. | This position is most commonly occupied by the isosteric Gln/Glu or by another acidic residue (Asp), but polar residues such as Asn and His have also been reported. A fully exposed, bulkier Arg side chain might have at most a small structural and/or functional impact. | – |
| c.1898G>A / p.Arg633Lys (SNP rs1198925225) | Not reported in ClinVar. Probably benign. | Conservative replacement of an almost strictly conserved, basic arginine by the lysine most common in mammalian MR sequences. Unlikely to have a global impact on the domain structure but might have a small effect on the conformation of the L1-3 loop and thus on receptor dimerization. | – |
| c.1903A>T / p.Thr635Ser (SNP rs745665712) | Not reported in ClinVar. Likely benign. | Conservative replacement of threonine by the serine found in several GR sequences, including some from primates. Unlikely to have any significant impact on the protein structure and function. | – |
| c.1921G>A / p.Asp641Asn (SNP rs756361771) | Not reported in ClinVar. Probably benign. | This solvent-exposed position is almost invariably occupied by an acidic residue, both in GR and MR, although the isosteric Asn and Ser have been reported in GR sequences from fishes. The replacement might have a minor impact on receptor multimerization (see below). | – |
| **c.1922A>T / p.Asp641Val (SNP rs104893908)** | Index patient: 58 y/o male homozygous for the mutation presented with severe hypertension and | Reduced affinity for DEX (3x) and transcriptional activity. Leads to enhanced multimerization at the DNA array. Modulated only two-thirds of WT-GR responsive genes. | [1,17–19] |

| | | | |
|---|---|---|---|
| | hypokalemia. No adrenal hyperplasia. Elevated ACTH. Both his son and nephew heterozygous for the mutation were asymptomatic and showed milder GC resistance. | This solvent-exposed position is invariably occupied by acidic residue or, less commonly, polar residues, both in GR and MR. Replacement by an aliphatic residue stabilizes transcriptionally impaired oligomers on chromatin. | |
| c.1933C>T / p.His645Tyr (SNP rs1816988536) | Not reported in ClinVar. Probably benign. | This position is not conserved: polar (mostly His/Gln, in mammals, birds and ray-finned fishes), basic (Arg, in reptiles) and acidic residues (Asp/Glu, in amphibians) have been reported. The side chain-less Gly is most common in cartilaginous fishes. An aromatic Tyr is exceptionally found in the *Bombina bombina* GR. Unlikely to have an important impact on the protein structure and function. | – |
| c.1943A>T,G / p.Tyr648Phe, Cys (SNP rs1333998849) | Not reported in ClinVar. Likely / probably benign. | Conservative replacement of a non-conserved aromatic residue by the Phe found in many GR sequences, including several from primates, or by a cysteine. Other residues commonly found at this position are aliphatic Leu/Met -almost strictly conserved in birds and reptiles-, and basic Lys/Arg residues in amphibians and ray-finned fishes. Unlikely to have an important impact on the protein structure and function. | – |
| c.1960C>T / p.His654Tyr (SNP rs767750427) | Not reported in ClinVar. Probably benign. | This position is not conserved, and reported sequences feature aliphatic (Ala/Val), polar (Ser, Thr, Asn, Gln, His) and basic residues (Lys/Arg). Might have a small impact on the DBD-LBD interaction. | – |
| c.1961A>G / p.His654Arg (SNP rs755126916) | Not reported in ClinVar. Probably benign. | Replaces a non-conserved residue by the arginine found in some GR sequences (see above). Might have a small impact on the DBD-LBD interaction. | – |
| c.1972G>A / p.Val658Ile (SNP rs113100205)[8] | Not reported in ClinVar. Probably benign. | Conservative replacement of a buried aliphatic residue by the Ile found in several amphibian GR sequences and conserved in amphibian MR. Unlikely to have any important structural or functional impact. | – |
| c.2011C>G / p.Leu671Val (SNP rs1816979477)[9] | Not reported in ClinVar. Probably pathogenic. | Conservative replacement of a buried aliphatic residue almost strictly conserved in both GR and MR by a less bulky Val. Might have a definite impact on the domain stability. | – |
| **c.2015T>C / p.Leu672Pro** | <u>Patient</u>: 60 y/o male heterozygous for the mutations presented with bilateral adrenal hyperplasia. No obvious clinical sign of Cushing's syndrome. Neither hypertension nor hypokalemia. | More sensitive to protein degradation than WT-GR. Unable to bind DEX. Remained exclusively cytoplasmic. Lacks transcriptional activity.<br>The non-conservative replacement of an almost strictly conserved, buried aliphatic residue might impair domain folding and stability, in line with the increased proteasomal degradation of this mutant. | [20] |

[8] Not validated.
[9] Not validated.

| | | | |
|---|---|---|---|
| c.2030A>G / p.Lys677Arg (SNP rs1813374572) | Not reported in ClinVar. Probably benign. | Conservative replacement of an almost strictly conserved, exposed basic residue. Lys is also much more common in MR sequences, although Arg is also reported. Unlikely to affect the domain structure but might have a minor impact on the protein function. | – |
| c.2034C>A,G / p.Asp678Glu (SNP rs258751)[10] | Polymorphism is significantly associated with steroid resistance in patients with primary nephrotic syndrome. (The A allele is linked to a reduced incidence of steroid resistance compared with the G allele). | Conservative replacement of a fully exposed acidic residue by the glutamate most common in mammals, and strictly conserved in birds and reptiles. The position is also occupied by Asp/Glu in MR sequences. Unlikely to have any direct impact on the protein structure and function. | [21,22] |
| **c.2035G>A / p.Gly679Ser (SNP rs104893914)** | Patient: 31 y/o female with hirsutism carrying the mutation in heterozygosity. Increased secretion of diurnal urinary cortisol and aldosterone. Normal blood pressure. | Reduced affinity for DEX (2x). Delayed nuclear translocation (2.5x). Preserved ability to bind to GREs and to interact with NCOA2 (only through AF-1). Decreased transcriptional activity in response to DEX (2x), but full capacity to repress TNF□-induced NF-□B activity. Conflicting results regarding the dominant negative effect on the WT receptor.<br>Replaces a strictly conserved, solvent exposed Gly, also conserved in other oxosteroid receptors, by a polar serine. Might affect domain folding and/or stability. | [23–25] |
| c.2036G>A / p.Gly679Asp (SNP rs1599646604) | Not reported in ClinVar, but likely pathogenic. | Replaces a strictly conserved, solvent exposed Gly by an acidic Asp. Might affect domain folding and/or stability (see above). | – |
| c.2050G>C / p.Glu684Gln (SNP rs1813368708) | Not reported in ClinVar. Likely benign. | Replaces a fully exposed, not conserved acidic residue by the isosteric Gln. In addition to the Glu most common in mammals, Gly, Ala, Pro, Ser, Thr and Asp are also found at this position. Unlikely to have any impact on the protein structure and function. | – |
| c.2059G>C / p.Asp687His (SNP rs762197361) | Not reported in ClinVar. Probably pathogenic. | This position is almost invariably occupied by an acidic residue in both GR and MR. Non-conservative replacement of this fully exposed residue by a His might have some functional impact. | – |
| c.2061T>G / p.Asp687Glu (SNP rs68012717)[11] | Not reported in ClinVar. Slightly increased levels of protein compared with WT, and increased GR□ receptor number. Likely benign. | Conservative replacement of a fully exposed aspartate by the glutamate strictly conserved in avian sequences and quite common also in reptiles and amphibians. Unlikely to have any impact on the protein structure and function. | [26] |
| c.2065A>C / p.Ile689Leu (SNP rs764474909) | Not reported in ClinVar. Likely benign. | Conservative replacement of a buried, strictly conserved aliphatic residue by the leucine found in the AR. Met strictly conserved at this position in MR sequences. Unlikely to have an important | – |

[10] Common polymorphism, in particular in African and Asian populations.
[11] Appears to be more common in Asian populations.

| | | | |
|---|---|---|---|
| | | impact on the protein structure and function. | |
| c.2090T>A / p.Leu697Gln (SNP rs1292760854) | Not reported in ClinVar, but likely pathogenic. | Replaces a buried aliphatic leucine strictly conserved in both GR and MR by the isosteric, polar Gln. Likely to impair folding and/or significantly decrease the stability of the LBD. | – |
| c.2097A>C / p.Lys699Asn (SNP rs1483819721) | Not reported in ClinVar. Probably pathogenic. | This position is almost invariably occupied by a lysine, and exceptionally by an Arg. The same residues are also invariably found in the MR. Replacement of this fully exposed basic residue by a polar Asn might have some structural or, more likely, functional impact. | – |
| c.2098G>A / p.Ala700Thr (SNP rs1406643989) | Not reported in ClinVar. Probably pathogenic. | Replacement of this strictly conserved, buried Ala by a bulkier polar Thr might result in steric clashes with H8 side chains (Glu661, Glu662). Might have some impact on domain stability. | – |
| c.2102T>C / p.Ile701Thr (SNP rs1274937680) | Not reported in ClinVar. Probably pathogenic. | Replacement of a strictly conserved aliphatic residue buried at the H9-H10 interface by a polar Thr might result in some destabilization of the local structure. | – |
| c.2107A>G / p.Lys703Glu (SNP rs760310145) | Not reported in ClinVar. Probably pathogenic. | Reversal-of-charge mutation of this exposed, almost strictly conserved residue eliminates a SUMOylation site, with likely functional consequences. | [27,28] |
| c.2109G>C / p.Lys703Asn (SNP rs529320390) | Not reported in ClinVar. Probably pathogenic. | Replacement of the almost strictly conserved, exposed lysine by a polar Asn. Gln is reported at this position in a few entries. Eliminates a SUMOylation site, with likely functional consequences. | [27,28] |
| c.2122T>C / p.Ser708Pro (SNP rs1282884786) | Not reported in ClinVar. Probably benign. | Although serine is strictly conserved at this position from amphibians to humans, Pro is found in several GR sequences from ray-finned fishes. (Along with Ala and Thr). Might have a minor impact on the structure of the L9-10 loop. | – |
| c.2126G>A,T / p.Ser709Asn,Ile (SNP rs1470677560) | Not reported in ClinVar. Probably pathogenic. | This position is almost invariably occupied by a serine. Replacement by either a bulkier polar residue (Asn) or by an aliphatic Ile might affect the conformations of the L9-10 loop. | – |
| c.2128C>A / p.Gln710Lys (SNP rs1813354187) | Not reported in ClinVar. Probably pathogenic. | Glutamine is strictly conserved at this position, both in the GR and the MR. Replacement by a basic Lys would impair the closed conformation of the L9-10 loop. | – |
| c.2130G>C / p.Gln710His (SNP rs1251767755) | Not reported in ClinVar. Probably pathogenic. | Replaces the strictly conserved Gln by a histidine. Might impair the closed conformation of the L9-10 loop. | – |
| c.2132A>G / p.Asn711Ser (SNP rs1813353161) | Not reported in ClinVar. Probably pathogenic. | Replacement of a strictly conserved asparagine by the less bulky Ser strictly conserved in MR from reptiles to humans. Might have a destabilizing effect on the closed conformation of the L9-10 loop. | – |
| c.2140C>T / p.Arg714Trp (SNP rs771583164) | Not reported in ClinVar, but likely pathogenic. | Replacement of this buried arginine strictly conserved in both GR and MR by a Trp would result in the loss of important stabilizing interactions between H10 and neighboring regions of the LBD. (Salt bridge to the Glu662 | – |

| | | | |
|---|---|---|---|
| | | carboxylate, H-bonds to main chain O atoms of Leu665). (See also below). | |
| **c.2141G>A /<br>p.Arg714Gln** | Patient #1: 2 y/o female carrying a nonsense mutation in the second allele (p.E198X) presented with vomiting, diarrhea and generalized seizure associated with severe hypertension, hypoglycemia and hypokalemia. She also had premature pubarche. Elevated levels of serum cortisol and plasma ACTH. Multiple small hemorrhagic infarcts detected by MRI at the age of 11 yrs. Computed tomography of the adrenals revealed bilateral adrenal hyperplasia.<br>Patient #2: 31 y/o female carrying the mutation in heterozygosity presented with infertility. Elevated serum and salivary cortisol. Slightly elevated or normal ACTH. Her 35-year-old sister carries the same mutation but is clinically healthy and has normal steroid hormone levels. | Reduced affinity for DEX (2x). Delayed nuclear translocation. Abnormal NCOA2 interaction and significantly reduced transactivation activity, with a dominant negative effect on the WT receptor. Loss of important stabilizing interactions between H10 and neighboring regions of the LBD (see above). Protein modeling suggests that the mutation transmits a conformational change to the LBP and the AF-2, resulting in a decreased binding affinity to ligand and to coactivators featuring the LXXLL motif. | [29–31] |
| c.2141G>T /<br>p.Arg714Leu<br>(SNP rs759445158) | Not reported in ClinVar, but likely pathogenic. | Loss of important stabilizing interactions between H10 and neighboring regions of the LBD (see above). | – |
| c.2151A>C /<br>p.Gln717His<br>(SNP rs770730144) | Not reported in ClinVar. Probably benign. | Replacement of an almost strictly conserved Gln by another polar residue rarely found in GR and MR. Might have a small impact on the local H10 structure. | – |
| **c.2177A>G /<br>p.His726Arg** | Patient: 30-year female carrying the mutation in heterozygosis presented with a long-standing story of hirsutism, acne, alopecia, anxiety, fatigue, and irregular menstrual cycles. Elevated plasma ACTH levels. | Decreased affinity for DEX (2x). Delayed nuclear translocation (4x). Abnormal NCOA2 interaction. Reduced transactivation (40%) and transrepression of NF-□B (30%). Does not exert a dominant-negative effect on WT GR□.<br>This position is invariably occupied by either His (from amphibians to humans and in cartilaginous fishes) or Gln (in several sequences from ray-finned fishes). Histidine is also much more common in MR sequences. The bulkier, positively charged Arg726 side chain might impair domain folding and/or stability. Modeling studies suggest reduced flexibility / increased rigidity of helix H10 due to the formation of stronger H-bonds. | [32] |

| | | | |
|---|---|---|---|
| c.2177A>C / p.His726Pro (SNP rs1813347977) | Not reported in ClinVar, but likely pathogenic. | Non-conservative replacement of this well conserved histidine by the helix breaker proline would lead to a disruption of helix H10, compromising domain folding and stability. | – |
| **c.2185G>A / p.Val729Ile (SNP rs1027058734)**[12] | Patient: 6 y/o male carrying the mutation in homozygosis presented with precocious pseudopuberty resulting from hyperandrogenism. High levels of serum cortisol and androgenic hormones. Her mother was heterozygous for the mutation. | Reduced affinity for DEX (2x) and cortisol (3x) and reduced transcriptional activity (4x).<br>This buried position is most commonly occupied by an aliphatic Val in both GR and MR, but the less bulky Ala and Gly have been reported in several GR sequences. Gly is also found in a few MR entries. Although conservative, the introduction of a bulkier Ile would have a definite impact on LBD folding and stability. (Clashes with Phe602 or Leu733). | [33] |
| c.2186T>C / p.Val729Ala (SNP rs1353048513) | Not reported in ClinVar. Likely benign. | Replaces a buried aliphatic valine by the Ala found in several GR sequences. Might have at most a minor impact on domain stability. | – |
| c.2189A>G / p.Glu730Gly (SNP rs1458571496) | Not reported in ClinVar. Likely benign. | Although an acidic glutamate is most common at his position, Gly has been reported in several GR sequences from ray-finned fishes and is most common in MR from amphibians and fishes. Might have at most a minor impact on the protein structure and function. | – |
| c.2193T>G / p.Asn731Lys (SNP rs1364678058) | Not reported in ClinVar. Probably benign. | Asn is most common at this position, but Ser is also repeatedly found, and Gly is almost strictly conserved in fishes. Replacement of this fully exposed residue by a basic lysine exceptionally reported in *Choloepus didactylus* GR is unlikely to have a large impact on the protein structure and function. | – |
| c.2198T>G / p.Leu733Arg (SNP rs376217889) | Not reported in ClinVar. Probably pathogenic. | Replaces a buried Leu strictly conserved in both GR and MR by a basic residue. Although the bulkier Arg side chain could be accommodated in the protein core in an extended conformation, donating H-bonds to the carbonyl O atom of Tyr598, this replacement is likely to have an impact on domain folding and stability. | – |
| c.2201A>G,C / p.Asn734Ser,Thr (SNP rs1229688908) | Not reported in ClinVar. Likely benign. | Replaces an exposed Asn by other polar residues commonly found in GR sequences, even from primates. Unlikely to have any impact on the protein structure and function. | – |
| c.2204A>G / p.Tyr735Cys (SNP rs780355678) | Not reported in ClinVar. Probably pathogenic. | Might lead to slightly reduced affinity for DEX (up to 2x), impaired interactions with NCOA1, and enhanced interactions with the Leu-Xxx-Xxx-Ile-Ile motif in NCOR1. (CORNR box 3; by similarity to tested mutations, Tyr735Phe/Val/Ser). This would result in impaired transactivating activity.<br>The position is almost invariably occupied by an aromatic Phe/Tyr from amphibians to humans, but Ile is also found in several fish sequences. Replacement by the less bulky Cys would selectively affect GC binding to the LBD and | [34,35] |

[12] Nucleotide 2,317 in the original publication.

| | | | |
|---|---|---|---|
| | | might have a minor effect on the local domain stability. (Loss of H-bond to the Asp641 carboxylate and of VdW contacts with Gln672). | |
| **c.2209T>C / p.Phe737Leu (SNP rs121909727)**[13] | Patient: 7 y/o male carrying the mutation in heterozygosis presented with severe hypertension, hypokalemia, and bilateral adrenal hyperplasia. Elevated ACTH. | Reduced affinity for DEX (2x). Delayed nuclear translocation (12x) but preserved ability to bind DNA. Abnormal interaction with NCOA2. Reduced transcriptional activity (2x). Exerted a dominant-negative effect on WT hGR only upon short-time exposure to the ligand.<br>Replacement of a Phe almost strictly conserved in both GR and MR by an aliphatic leucine might affect LBD folding and/or stability. | [36] |
| c.2223G>T / p.Leu741Phe (SNP rs745976068) | Not reported in ClinVar. Probably pathogenic. | This position is almost invariably occupied by an aliphatic residue (Leu from reptiles to humans, Val most common in fishes), but Thr has also been reported in several amphibian sequences. Replacement of this well exposed residue by a bulkier Phe is unlikely to affect protein structure but might enhance receptor oligomerization. | – |
| c.2225A>T / p.Asp742Val (SNP rs1445952313) | Not reported in ClinVar. Probably pathogenic. | This position is invariably occupied by an acidic Asp (from amphibians to humans), the isosteric Asn (in ray-finned fishes) or Glu/Gln (in cartilaginous fishes), and is strictly conserved as Glu in MR. Presence of an aliphatic residue at this exposed position might enhance non-productive multimerization, similar to Asp641Val. | – |
| c.2227A>G / p.Lys743Glu (SNP rs1247900918) | Not reported in ClinVar. Probably pathogenic. | Reversal-of-charge replacement of a fully exposed, strictly conserved residue. Generates a strong electronegative potential with the carboxylates of nearby residues (Asp742, Glu748, and Glu751). Might impair domain folding and/or stability. | – |
| c.2228A>T / p.Lys743Met (SNP rs2151472980)[14] | Not reported in ClinVar. Probably pathogenic. | Replaces strictly conserved Lys by an aliphatic Met. Unlikely to affect domain folding but might increase non-productive oligomerization. | – |
| c.2231C>A / p.Thr744Asn (SNP rs778847361) | Not reported in ClinVar. Likely benign. | Replaces an exposed Thr by another polar residue found in several mammalian GR sequences. Unlikely to have a significant impact on the protein structure and function. | – |
| c.2233A>G / p.Met745Val (SNP rs757831210) | Not reported in ClinVar. Probably benign. | Although an aliphatic Met/Leu is much more common at this position, the polar Thr is also found in several avian GR sequences. Replacement by the less bulky Val would impair VdW interactions with nearby residues (Thr739, Asp742), with a small impact on the local protein structure. | – |
| c.2237G>T / p.Ser746Ile (SNP rs866664218) | Not reported in ClinVar. Probably pathogenic. | A polar serine is much more common at this position, but the isosteric Cys, another polar residue, Asn, and the basic Arg have also been reported. Introduction of an aliphatic side chain at this exposed position might enhance the tendency to non-productive oligomerization. | – |

[13] Not validated.
[14] Not validated.

| | | | |
|---|---|---|---|
| **c.2241T>G / p.Ile747Met (SNP rs104893910)** | Patient: 18-year female carrying the mutation in heterozygosis presented with hyperandrogenism, cystic acne and hirsutism at puberty, and persistent oligo-amenorrhea. Elevated plasma cortisol concentration. Her father and her two brothers, also heterozygous for the mutation, were phenotypically silent despite elevated levels of cortisol. | Slight delay in nuclear translocation. Reduced affinity for DEX (2x). Interacts with NCOA2 through AF-1 but not through AF-2. Lower transcriptional activity (4- to 5-fold at low DEX concentrations, 2x at high concentrations). Dominant negative inhibitor of WT GR-induced transactivation. This buried position is invariably occupied by either Ile or Val, also in other oxosteroid receptors. Replacement by Met would seem to be tolerated but might differentially affect GC binding. Might lead to enhanced interactions with the N-terminus of H3 (Trp557, Met560). | [37] |
| c.2242G>A / p.Glu748Lys (SNP rs1813187040) | Not reported in ClinVar. Probably pathogenic. | Reversal-of-charge mutation of a fully exposed, strictly conserved acidic residue. Position also conserved as Asp/Glu in other oxosteroid receptors. Might have some impact on PPIs, and in particular on the binding of coregulators. | – |
| c.2251G>A / p.Glu751Lys (SNP rs1208097433) | Not reported in ClinVar. Probably pathogenic. | Reversal-of-charge mutation of a fully exposed, almost strictly conserved acidic residue. Might affect secondary interactions with coregulators. | – |
| **c.2259A>T / p.Leu753Phe (SNP rs121909726)[15]** | GC-resistant mutant isolated from GC-sensitive human leukemic CEM-C7 cells. | Reduced affinity for DEX (5x) and transactivating activity (8x). Replaces buried Leu strictly conserved in both GR and MR by an aromatic Phe. Unlikely to impair domain folding but would directly or indirectly affect hormone binding. Might affect interactions with H3 residues (Gly568, Val571) and with Trp600. | [38,39] |
| c.2264A>T / p.Glu755Val (SNP rs1219069553) | Not reported in ClinVar, but likely pathogenic. | Replaces strictly conserved, acidic AF-2 residue by an aliphatic valine, with a large impact on coregulator binding. | – |
| **c.2269A>G / p.Ile757Val** | Patient: 23 y/o female carrying the mutation in heterozygosis. | Unable to bind DEX and to translocate into the nucleus. Conservatively replaces a buried aliphatic residue strictly conserved in both GR and MR by a less bulky valine. Might decrease the stability of the LBD. | [40] |
| c.2272A>T / p.Thr758Ser (SNP rs1437232377) | Not reported in ClinVar. Likely benign. | Conservative replacement of Thr by the serine almost strictly conserved in GR sequences from avians, reptiles and amphibians, and also most common in the MR. Unlikely to have any impact on the protein structure and function. | – |
| c.2275A>G / p.Asn759Asp (SNP rs1813181716) | Not reported in ClinVar. Probably benign. | Replaces strictly conserved Asn by the isosteric Asp strictly conserved in the MR. Might have some impact on the binding of specific coregulators. | – |
| c.2276A>G / p.Asn759Ser (SNP rs756497284) | Not reported in ClinVar. Probably benign. | Replaces strictly conserved Asn by a less bulky, polar serine. Could have a small impact on the binding of some coregulators. | – |
| c.2281A>G / p.Ile761Val | Not reported in ClinVar. Probably benign. | This position is invariably occupied by an aliphatic Ile/Leu in both GR and MR. Although the | – |

[15] Not validated.

| | | | |
|---|---|---|---|
| (SNP rs1813180628) | | replacement is conservative, the less bulky Val residue lacks VdW contacts with nearby side chains (Ile757), which might have a small impact on the domain stability. | |
| c.2282T>C / p.Ile761Thr (SNP rs2151472803)[16] | Not reported in ClinVar. Probably benign. | This position is invariably occupied by an aliphatic Ile/Leu in both GR and MR. The presence of a polar residue at this position might weaken interactions with the side chains of nearby residues (see above). Likely to have a small impact on the domain folding and stability. | – |
| c.2284C>A / p.Pro762Thr (SNP rs1813179596) | Not reported in ClinVar. Probably pathogenic. | This proline residue is almost strictly conserved (Ser exceptionally reported in the *Elephantulus edwardii* sequence) and is also conserved in other oxosteroid receptors. Replacement by a polar Thr might affect local protein folding. | – |
| c.2288A>G / p.Lys763Arg (SNP rs1813179056) | Not reported in ClinVar. Likely benign. | Conservative replacement of a fully exposed, almost strictly conserved basic residue by the Arg exceptionally found in a couple of GR sequences. Lys is also the most common in MR, but there are reversal-of-charge replacements to Asp/Glu in MR sequences from cartilaginous fishes. Unlikely to have an important impact on the protein structure and function. | – |
| c.2294C>T / p.Ser765Leu (SNP rs1318675518) | Not reported in ClinVar. Probably pathogenic. | This fully exposed position is invariably occupied by a polar Ser -or, less frequently, Thr- from amphibians to humans, and by a basic lysine in ray-finned fishes. Met is strictly conserved in cartilaginous fishes. Replacement by an aliphatic leucine is unlikely to have any impact on the domain folding but might enhance non-productive multimerization. | – |
| c.2296A>G / p.Asn766Asp (SNP rs186936077) | Not reported in ClinVar. Likely benign. | Replacement of a non-conserved Asn by the isosteric Asp most commonly found in ray-finned fishes. Unlikely to have any significant impact on the protein structure and function. | – |
| c.2299G>A / p.Gly767Arg (SNP rs1813176820) | Not reported in ClinVar. Probably pathogenic. | Replaces strictly conserved glycine, also conserved in other oxosteroid receptors, by a bulky, basic Arg. Increases electropositive character of the surface epitope created by nearby lysine side chains at positions 677, 770, 771, and 777. Might impair folding of the F domain. | – |
| c.2302A>G / p.Asn768Asp (SNP rs754526334) | Not reported in ClinVar. Probably benign. | Isosteric replacement of the asparagine almost strictly conserved from amphibians to humans. Ser is much more common in ray-finned fishes, while Met is conserved in cartilaginous fishes. Might have a minor impact on the local protein conformation due to electrostatic repulsion from the Asp678 carboxylate. | – |
| c.2306T>C / p.Ile769Thr (SNP rs1412555345) | Not reported in ClinVar. Likely benign. | This position is invariably occupied by an aliphatic Ile (most common from amphibians to humans), Val (in ray-finned fishes) or Ala (in cartilaginous fishes). Thr is strictly conserved in MR sequences from fishes. The presence of this polar residue | – |

[16] Not validated.

| | | | |
|---|---|---|---|
| | | might have a minor impact on the domain folding and/or stability. | |
| c.2307C>G / p.Ile769Met (SNP rs886060060)[17] | Not reported in ClinVar. Likely benign. | Conservative replacement of an aliphatic but not conserved residue (see above). Might have at most a minor impact on the protein structure and function. | – |
| c.2309A>G / p.Lys770Arg (SNP rs751136795) | Not reported in ClinVar. Likely benign. | Conservative replacement of a basic lysine by an Arg most common in GR sequences from amphibians. Unlikely to have a significant impact on the protein structure and function. | – |
| c.2317C>A / p.Leu773Met (SNP rs765982151) | Not reported in ClinVar. Likely benign. | Conservative replacement of a partially buried aliphatic residue strictly conserved from ray-finned fishes to humans. Arg strictly conserved in cartilaginous fishes. Might have a small impact on domain folding and/or stability. | – |
| **c.2318T>C / p.Leu773Pro (SNP rs104893912)** | Patient: 29 y/o female carrying the mutation in heterozygosis presented with a long-standing history of fatigue, anxiety, menstrual irregularities, hyperandrogenism, and hypertension. Acne, hirsutism. Elevated concentrations of serum cortisol, plasma ACTH and androgenic hormones. | There are no overt effects on the mutant protein, but it is more dynamic with slightly lower thermal stability. Reduced affinity for DEX (2.4-2.6x). Delayed nuclear translocation (2.5x). Preserved ability to bind to DNA, although with altered dimerization behavior. Interacts with NCOA2 only through AF-1. Decreased transcriptional activity (2x). Dominant negative effect on the WT receptor. The non-conservative replacement might affect the conformation of the F domain. | [41,42] |
| c.2323C>T / p.His775Tyr (SNP rs773260276) | Not reported in ClinVar. Likely benign. | Replaces histidine strictly conserved in both GR and MR by an aromatic tyrosine. Unlikely to have an important impact on the overall domain structure and might actually increase domain stability through VdW interactions with Tyr716. | – |

### References to Supplementary Table 2.


1. Jiménez-Panizo A, Alegre-Martí A, Tettey TT, Fettweis G, Abella M, Antón R, et al. The multivalency of the glucocorticoid receptor ligand-binding domain explains its manifold physiological activities. Nucleic Acids Res 2022;50(22):13063–82.

2. Ma L, Tan X, Li J, Long Y, Xiao Z, De J, et al. A novel glucocorticoid receptor mutation in primary generalized glucocorticoid resistance disease. Endocr Pract 2020;26(6):651–9.

3. Zhu HJ, Dai YF, Wang O, Li M, Lu L, Zhao WG, et al. Generalized glucocorticoid resistance accompanied with an adrenocortical adenoma and caused by a novel point mutation of human glucocorticoid receptor gene. Chin Med J (Engl) 2011;124(4):551–5.

4. Nicolaides NC, Skyrla E, Vlachakis D, Psarra AG, Moutsatsou P, Sertedaki A, et al. Functional characterization of the hGRαT556I causing Chrousos syndrome. Eur J Clin Invest 2016;46(1):42–9.

5. Karl M, Lamberts SW, Koper JW, Katz DA, Huizenga NE, Kino T, et al. Cushing's disease preceded by


[17] Not validated.

generalized glucocorticoid resistance: clinical consequences of a novel, dominant-negative glucocorticoid receptor mutation. Proc Assoc Am Physicians 1996;108(4):296–307.

6. Kino T, Stauber RH, Resau JH, Pavlakis GN, Chrousos GP. Pathologic human GR mutant has a transdominant negative effect on the wild-type GR by inhibiting its translocation into the nucleus: Importance of the ligand-binding domain for intracellular GR trafficking. J Clin Endocrinol Metab 2001;86(11):5600–8.
7. Panek M, Pietras T, Antczak A, Fabijan A, Przemêcka M, Górski P, et al. The N363S and I559N single nucleotide polymorphisms of the h-GR/NR3C1 gene in patients with bronchial asthma. Int J Mol Med 2012;30(1):142–50.
8. Lind U, Greenidge P, Gillner M, Koehler KF, Wright A, Carlstedt-Duke J. Functional probing of the human glucocorticoid receptor steroid- interacting surface by site-directed mutagenesis: Gln-642 plays an important role in steroid recognition and binding. J Biol Chem 2000;275(25):19041–9.
9. Laulhé M, Kuhn E, Bouligand J, Amazit L, Perrot J, Lebigot E, et al. A novel mutation in the NR3C1 gene associated with reversible glucocorticoid resistance. Eur J Endocrinol 2024;190(4):284–95.
10. Mendonca BB, Leite M V, De Castro M, Kino T, Elias LLK, Bachega TAS, et al. Female pseudohermaphroditism caused by a novel homozygous missense mutation of the GR gene. J Clin Endocrinol Metab 2002;87(4):1805–9.
11. Lind U, Greenidge P, Gustafsson JÅ, Wright APH, Carlstedt-Duke J. Valine 571 functions as a regional organizer in programming the glucocorticoid receptor for differential binding of glucocorticoids and mineralocorticoids. J Biol Chem 1999;274(26):18515–23.
12. Nicolaides NC, Roberts ML, Kino T, Braatvedt G, Hurt DE, Katsantoni E, et al. A novel point mutation of the human glucocorticoid receptor gene causes primary generalized glucocorticoid resistance through impaired interaction with the LXXLL motif of the p160 coactivators: Dissociation of the transactivating and transreppressive activities. J Clin Endocrinol Metab 2014;99(5):902–7.
13. Cannavò S, Santarpia L, Karl M, Facchiano A, Marabotti A, Benvenga S, et al. A novel point mutation in the ligand-binding domain of the human glucocorticoid receptor (hGR) in a patient with glucocorticoid resistance. Int J Disabil Hum Dev 2007;6(1):105–12.
14. Seitz T, Thoma R, Schoch GA, Stihle M, Benz J, D'Arcy B, et al. Enhancing the stability and solubility of the glucocorticoid receptor ligand-binding domain by high-throughput library screening. J Mol Biol 2010;403(4):562–77.
15. Murani E, Reyer H, Ponsuksili S, Fritschka S, Wimmers K. A substitution in the ligand binding domain of the porcine glucocorticoid receptor affects activity of the adrenal gland. PLoS One 2012;7(9):e45518.
16. Reyer H, Ponsuksili S, Kanitz E, Pöhland R, Wimmers K, Murani E. A natural mutation in helix 5 of the ligand binding domain of glucocorticoid receptor enhances receptor-ligand interaction. PLoS One 2016;11(10):e0164628.
17. Chrousos GP, Vingerhoeds A, Brandon D, Eil C, Pugeat M, DeVroede M, et al. Primary cortisol resistance in man. A glucocorticoid receptor-mediated disease. J Clin Invest 1982;69(6):1261–9.
18. Chrousos GP, Vingerhoeds ACM, Loriaux DL, Lipsett MB. Primary cortisol resistance: A family study. J Clin Endocrinol Metab 1983;56(6):1243–5.
19. Hurley DM, Accili D, Stratakis CA, Karl M, Vamvakopoulos N, Rorer E, et al. Point mutation causing a

single amino acid substitution in the hormone binding domain of the glucocorticoid receptor in familial glucocorticoid resistance. J Clin Invest 1991;87(2):680–6.

20. Vitellius G, Fagart J, Delemer B, Amazit L, Ramos N, Bouligand J, et al. Three novel heterozygous point mutations of NR3C1 causing glucocorticoid resistance. Hum Mutat 2016;37(8):794–803.
21. Liu J, Wan Z, Song Q, Li Z, He Y, Tang Y, et al. NR3C1 gene polymorphisms are associated with steroid resistance in patients with primary nephrotic syndrome. Pharmacogenomics 2018;19(1):45–60.
22. Wu C, Fang F, Zhan X, Wei Y. The association between glucocorticoid receptor (NR3C1) gene polymorphism and difficult-to-treat rhinosinusitis. Eur Arch Oto-Rhino-Laryngology 2022;279(8):3981–7.
23. Ruiz M, Lind U, Gåfvels M, Eggertsen G, Carlstedt-Duke J, Nilsson L, et al. Characterization of two novel mutations in the glucocorticoid receptor gene in patients with primary cortisol resistance. Clin Endocrinol (Oxf) 2001;55(3):363–71.
24. Charmandari E, Kino T, Ichijo T, Zachman K, Alatsatianos A, Chrousos GP. Functional characterization of the natural human glucocorticoid receptor (hGR) mutants hGRαR477H and hGRαG679S associated with generalized glucocorticoid resistance. J Clin Endocrinol Metab 2006;91(4):1535–43.
25. Ruiz M, Hedman E, Gåfvels M, Eggertsen G, Werner S, Wahrenberg H, et al. Further characterization of human glucocorticoid receptor mutants, R477H and G679S, associated with primary generalized glucocorticoid resistance. Scand J Clin Lab Invest 2013;73(3):203–7.
26. Niu N, Manickam V, Kalari KR, Moon I, Pelleymounter LL, Eckloff BW, et al. Human glucocorticoid receptor α gene (NR3C1) pharmacogenomics: Gene resequencing and functional genomics. J Clin Endocrinol Metab 2009;94(8):3072–84.
27. Tian S, Poukka H, Palvimo JJ, Jänne OA. Small ubiquitin-related modifier-1 (SUMO-1) modification of the glucocorticoid receptor. Biochem J 2002;367(3):907–11.
28. Druker J, Liberman AC, Antunica-Noguerol M, Gerez J, Paez-Pereda M, Rein T, et al. RSUME enhances glucocorticoid receptor SUMOylation and transcriptional activity. Mol Cell Biol 2013;33(11):2116–27.
29. Nader N, Bachrach BE, Hurt DE, Gajula S, Pittman A, Lescher R, et al. A novel point mutation in helix 10 of the human glucocorticoid receptor causes generalized glucocorticoid resistance by disrupting the structure of the ligand-binding domain. J Clin Endocrinol Metab 2010;95(5):2281–5.
30. Molnár Á, Patócs A, Likó I, Nyíro G, Rácz K, Tóth M, et al. An unexpected, mild phenotype of glucocorticoid resistance associated with glucocorticoid receptor gene mutation case report and review of the literature. BMC Med Genet 2018;19(1):37.
31. Tatsi C, Stratakis CA. The genetics of pituitary adenomas. J Clin Med. 2020;9(1):30.
32. Nicolaides NC, Geer EB, Vlachakis D, Roberts ML, Psarra AMG, Moutsatsou P, et al. A novel mutation of the hGR gene causing Chrousos syndrome. Eur J Clin Invest 2015;45(8):782–91.
33. Malchoff DM, Brufsky A, Reardon G, McDermott P, Javier EC, Bergh CH, et al. A mutation of the glucocorticoid receptor in primary cortisol resistance. J Clin Invest 1993;91(5):1918–25.
34. Stevens A, Garside H, Berry A, Waters C, White A, Ray D. Dissociation of steroid receptor coactivator 1 and nuclear receptor corepressor recruitment to the human glucocorticoid receptor by modification of

the ligand-receptor interface: The role of tyrosine 735. Mol Endocrinol 2003;17(5):845–59.

**Supplementary Table 3 Legend. Contribution of different energy terms to the stability of major protein-protein interfaces in the current GR-LBD crystal.** Energy contributions are given in kcal/mol. The total pseudoenergy is calculated as the sum of electrostatic, desolvation, and one-tenth of the Van-der-Waals (VdW) term. Note that formation of the Tyr545 homodimers is driven by electrostatic interaction, while the desolvation effect is more relevant at the Trp712 interface.

| Dimer | Electrostatic | Desolvation | VdW | Total |
|---|---|---|---|---|
| **Basic, Tyr545-centered dimers** | | | | |
| A·B | –15.91 | 8.86 | –54.28 | –12.48 |
| C·D | –16.51 | 10.07 | –54.15 | –11.86 |
| E·F | –15.37 | 14.30 | –52.11 | –6.29 |
| G·H | –13.79 | 12.18 | –54.97 | –7.11 |
| I·J | –15.74 | 7.79 | –54.19 | –13.38 |
| K·L | –11.18 | 14.18 | –57.05 | –2.71 |
| M·N | –15.30 | 13.72 | –59.95 | –7.57 |
| O·P | –12.90 | 11.90 | –48.0 | –5.80 |
| **Pro637-centered tetramers-of-dimers** | | | | |
| B:C | -2.96 | -6.51 | -46.57 | -14.13 |
| E:(B:C) | -10.10 | -9.00 | -69.93 | -26.09 |
| H:(B:C) | -9.38 | -1.83 | -43.79 | -15.59 |
| J:N | -2.73 | -10.86 | -24.78 | -16.07 |
| K:(J:N) | -9.57 | -9.46 | -65.09 | -25.53 |
| O:(J:N) | -14.82 | -0.64 | -65.57 | -22.02 |
| **Trp712-centered dimers-of-dimers** | | | | |
| A:P | –0.72 | –24.44 | –44.41 | –29.61 |
| C:M | –4.04 | –22.73 | –66.98 | –33.47 |
| D:L | –1.80 | –27.56 | –52.64 | –34.62 |
| E:I | 0.96 | –30.86 | –42.32 | –34.14 |
| G:K | –2.94 | –22.81 | –37.18 | –29.54 |
| H:N | 0.19 | –20.33 | –43.01 | –24.44 |

**Supplementary Table 4 Legend. Relationship between the different Trp712-centered dimers of dimers.**

The RMSDs between the two dimers are given in Å. Note the separation in four clusters, formed by dimers D:L / E:I, A:P / H:N, C:M, and G:K (color-coded according to Supplementary Figure 5).

| Dimer | D:L | E:I | A:P | H:N | C:M | G:K |
|---|---|---|---|---|---|---|
| **D:L** | 0.0 | 2.1 | 5.1 | 5.6 | 18.2 | 19.1 |
| **E:I** | 2.1 | 0.0 | 4.1 | 5.1 | 18.5 | 19.0 |
| **A:P** | 5.1 | 4.1 | 0.0 | 2.1 | 18.6 | 17.5 |
| **H:N** | 5.6 | 5.1 | 2.1 | 0.0 | 17.0 | 15.7 |
| **C:M** | 18.2 | 18.5 | 18.6 | 17.0 | 0.0 | 8.2 |
| **G:K** | 19.1 | 19.0 | 17.5 | 15.7 | 8.2 | 0.0 |

**Video S1 Legend. 3D model of DNA-bound multidomain GR, related to Figure 4.** The model was generated by superimposing the non-canonical, Tyr545-centered GR-LBD homodimer on the recently reported crystal structure, 7PRW. The unmodified monomer from 7PRW is shown as a cyan cartoon, while the one generated according to the Tyr545-centered dimer is colored salmon. Bound DEX molecules are given as color-coded spheres, and $Zn^{2+}$ ions in the DBD modules as purple spheres. The pair of stacked tyrosine residues (Tyr545/Tyr545') at the center of the dimer interface and the supporting residues of the L5-6 loop (Pro625, Ile628) are shown as color-coded sticks.

**Video S2 Legend. Arrangement of GR-LBD monomers in the Pro637 setting, related to Figures S4 and 5.** Formation of the ASU of the crystal is presented as a sequential assembly process, in which dimers A·B and C·D associate first through the L6-7 loops from monomers B and C featuring Leu636 / Pro637 residues. Next, dimers E·F and G·H dock asymmetrically onto the 'cradles' formed at the B:C interface, through the L1-3 and L6-7 loops of monomers E and H. The octamer is completed by contacts between the peripheral monomers, A-F and D-G. A similar pathway generates the second octamer in the ASU, formed by GR-LBDs I-P.

**Video S3 Legend. Interconversion between the six GR-LBD dimers-of-dimers formed in the Trp712 setting, related to Figure 5.**

For simplicity, one dimer is set as reference, and only the monomer from the 'moving' dimer that contacts the Trp712 interface is shown. Note the central position of Trp712/712', Phe715/715' and Tyr716/716' aromatic side chains at these interfaces. Note also the large conformational space covered by the second molecule, when the first monomer is held fixed. The DNA-bound DBD of this monomer is included to inform the viewer about the approximate position of chromatin.
